# Repressive gene regulation synchronizes development with cellular metabolism

**DOI:** 10.1101/548032

**Authors:** Justin J. Cassidy, Sebastian Bernasek, Rachael Bakker, Ritika Giri, Nicolás Peláez, Bryan Eder, Anna Bobrowska, Neda Bagheri, Luis A. Nunes Amaral, Richard W. Carthew

## Abstract

Metabolic conditions affect the developmental tempo of most animal species. Consequently, developmental gene regulatory networks (GRNs) must faithfully adjust their dynamics to a variable time scale. We find evidence that layered weak repression of genes provides the necessary coupling between GRN output and cellular metabolism. Using a mathematical model that replicates such a scenario, we find that lowering metabolism corrects developmental errors that otherwise occur when different layers of repression are lost. Through mutant analysis, we show that gene expression dynamics are unaffected by loss of repressors, but only when cellular metabolism is reduced. We further show that when metabolism is lowered, formation of a variety of sensory organs in *Drosophila* is normal despite loss of individual repressors of transcription, mRNA stability, and protein stability. We demonstrate the universality of this phenomenon by experimentally eliminating the entire microRNA family of repressors, and find that all microRNAs are rendered unnecessary when metabolism is reduced. Thus, layered weak repression provides robustness through error frequency suppression, and may provide an evolutionary route to a shorter reproductive cycle.

## INTRODUCTION

Animal development occurs over a defined timescale, which requires control of the rates of developmental processes. Developmental timescales are an intrinsic feature of a species, and are not necessarily determined by external clocks (Ebisuya and Briscoe, 2018). Rather, the pace of development is encoded in the genome. Development occurs via a stereotypic sequence of events involving cell division, growth, movement, apoptosis, polarization, and differentiation. Correct assembly of functional structures depends upon synchronization of cell division and differentiation events (Foe, 1989; Sulston et al., 1983). Small variation in timing produces variation in structure that is observed between individuals (Francesconi and Lehner, 2014; Poullet et al., 2016). Abnormal timing can result in structural defects that lead to compromised survival (Moss, 2007).

While developmental tempo is a fundamental property of a species, it can vary under different conditions. For example, temperature affects the pace of development in many ectotherms, such as arthropods, nematodes, fish, and reptiles (Atlas, 1935; Davidson, 1944; Kuntz and Eisen, 2014; Zuo et al., 2011). Diet and food intake also affect organismal growth rate and the pace of development for many species, including humans (Arendt, 1997; Brown et al., 2004; Metcalfe and Monaghan, 2001; Pontzer et al., 2016). Finally, cellular metabolism can alter the pace of development. For example, the evolutionarily conserved *Clk1* gene encodes a mitochondrial enzyme necessary for normal cellular respiration (Felkai et al., 1999), and loss of the *clk1* gene in nematodes and mice results in developmental delays (Levavasseur et al., 2001; Nakai et al., 2001; Wong et al., 1995). In *Drosophila*, restricting glucose consumption by cells slows development (Brogiolo et al., 2001; Layalle et al., 2008; Rulifson et al., 2002; Shingleton et al., 2005). West and colleagues formulated a general quantitative model that relates developmental tempo to both cellular metabolic rate and temperature (Gillooly et al., 2002). Strikingly, the model fits meta-data spanning several kingdoms, suggesting a universal relationship between metabolism and developmental tempo.

Many developmental processes involve specification of different cell types in a stereotyped sequence. All of these differentiated cell types originate from progenitor cells. The sequence of cell differentiation is driven by changes in the gene expression program within progenitors. Gene regulators, typically transcription factors, are sequentially activated and repressed, resulting in transient periods of increased activity. During these periods, they change gene expression in the progenitors. This coincides with and causes a temporal series of cell fate decisions. Since these regulators frequently interact with one another, the entire cascade constitutes a gene regulatory network (GRN). Such GRNs have been characterized for embryogenesis (Cusanovich et al., 2018; Davidson and Erwin, 2006; Lawrence, 1992), development of the central nervous system (Kohwi and Doe, 2013), and development of the sensory nervous system (Cepko, 2014). Because the tempo of development can vary, GRN dynamics must be able to reliably adjust to a variable timing mechanism. Therefore, understanding how these GRNs adapt to a variable timescale is crucial for understanding the mechanisms of animal development.

Phenomenological observations suggest that there are limits to the timescales to which development may adapt. While broiler chickens have been successfully bred for rapid growth, frequent abnormalities in musculoskeletal development are evident in such breeds (Julian, 2005; Whitehead et al., 2003). Animals (and humans) experience hyper-normal growth rates if they initially experience delayed growth (Arendt, 1997). Such compensatory growth is linked to a variety of developmental and physiological defects (Metcalfe and Monaghan, 2001). Conversely, slowing growth can alleviate defects caused by mutations that impair development. As first noted by T.H. Morgan, morphological phenotypes can be suppressed by limiting the nutrition of mutant animals (Child, 1939; Morgan, 1915; Morgan, 1929; Sang and Burnet, 1963). Likewise, raising animals under lowered temperatures can sometimes suppress the phenotypes of mutations that are not classical *ts* alleles (Child, 1935; Krafka, 1920; Lewis et al., 1980; Villee, 1943). Collectively, these observations suggest an unknown mechanism ensures successful developmental outcomes amidst variability in developmental tempo.

Here, we have explored this mechanism. We find that impairing gene repression in GRNs causes developmental errors but only when cell metabolism and growth rate are normal. When either energy metabolism or protein anabolism are reduced, developmental errors are reduced or even suppressed. We find that this relationship between metabolism and repression is so prevalent that the entire microRNA family becomes unnecessary when metabolism is slowed. Using a general quantitative modeling framework for regulated gene expression, we show that multiple layers of weak repression render gene expression dynamics independent of variable biochemical rates. When rates are modestly reduced, fewer repressors are needed to ensure normal expression dynamics. We experimentally validate this model prediction by following GRN dynamics in *Drosophila*. Our findings support a new mechanism whereby layers of gene repression allow development to occur over a wider range of time scales, enabling development to proceed faster if metabolic conditions allow for it. The need for flexible error frequency suppression could provide an evolutionary impetus for the high prevalence of genetic redundancy.

## RESULTS

Developmental patterns arise from directed dynamics of cell-cell signaling and gene regulation. The sensory organs of *Drosophila* are a classic system with which to study these phenomena (Quan and Hassan, 2005). A broad collection of gene mutations has specific effects on the formation of various sensory organs, and these mutations have been instrumental in uncovering the molecular mechanisms of sensory organ development. The affected genes encode transcription factors, microRNAs, signaling factors, and other gene regulators. We used such gene mutations to readdress the relationship between reduced metabolism and phenotype suppression that was first observed by Morgan (Morgan, 1915; Morgan, 1929). We did so by scoring *Drosophila* sensory mutant phenotypes under conditions of reduced energy metabolism. We generated animals that had reduced metabolism by genetic ablation of their insulin producing cells (IPCs) in the brain (Fig. 1A). This ablation reduces the amount of glucose that cells consume (Rulifson et al., 2002), resulting in 70% slower development (Fig. 1B), and small but normally proportioned adults (Fig. 1C) (Rulifson et al., 2002).

**Figure 1.**
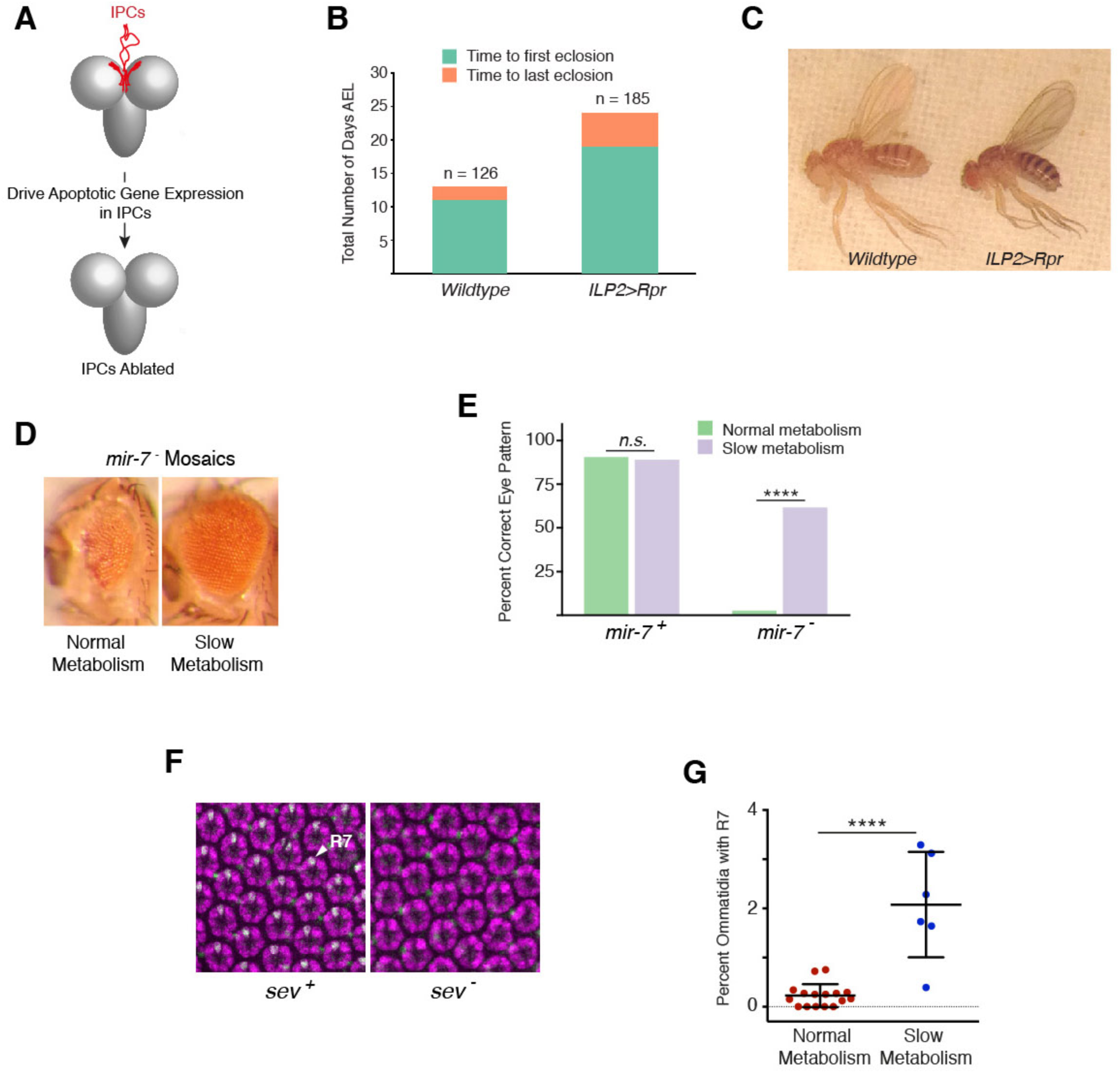
Eye developmental defects are rescued by slower energy metabolism. **(A)** Strategy to ablate IPCs (red) in the young fly brain. Gal4 expressed under control of the promoter for the Insulin-Like Peptide 2 (ILP2) gene drives production of the pro-apoptotic protein Reaper (Rpr) specifically in IPCs of the brain. **(B)** The number of days after egg laying (AEL) at which the first individual in either wildtype or *ILP2>Rpr* populations eclosed (hatched from pupa into adult) is shown, as is the time at which the last individual in each population eclosed. Population sizes for wildtype and *ILP2>Rpr* were 126 and 185, respectively. **(C)** Adult body size is affected by IPC ablation. Two females that were raised at the same time and temperature. The left y w animal has normal metabolism, whereas the right animal has slowed metabolism due to ablation of its IPCs. **(D)** Genetically mosaic individuals with a mir-7+ body and a mir-7 mutant eye. Left individual with mispatterned eye has its IPCs intact while the right individual with a normally patterned eye has had its IPCs ablated by *ILP2>Rpr*. **(E)** Eye patterning is more normal if mosaic individuals slowly metabolize due to IPC ablation. Sample population sizes were between 264 and 467 individuals. *P* values from Chi-square test with Yates’ correction. **(F)** Eye cells stained for specific protein markers such that R7 cells (white) can be distinguished from other R cells (purple) and bristle cells (green). Each ring-like cluster of R cells is an ommatidium. Null mutation of sev results in no R7 cells (right). **(G)** Slow metabolism due to IPC ablation increases the fraction of ommatidia that contain an R7 cell in sev mutants. Each data point represents one eye sample; between 481 and 837 ommatidia were scored for R7 cells within each eye sample. P value is from a one-way ANOVA with Bonferroni correction.

### Mutation of Repressors Has Less Impact When Metabolism is Reduced

We first examined mutations affecting formation of the compound eye. The microRNA miR-7 represses expression of the Yan transcription factor in the developing eye (Li and Carthew, 2005). Yan protein is transiently expressed in the eye (Pelaez et al., 2015), and is cleared from differentiating photoreceptor (R) cells by multiple repressors acting on its transcription, mRNA stability, and protein stability (Graham et al., 2010). When the *mir-7* gene was specifically ablated in the compound eye of an otherwise wildtype animal, it resulted in small malformed adult eyes due to errors in R cell differentiation (Fig. 1D). This phenotype was highly penetrant in genetically mosaic animals (Fig. 1E). However, when energy metabolism was slowed by IPC ablation, loss of *mir-7* was much less important for the formation of correctly patterned eyes (Fig. 1E). We also examined mutations affecting cell-cell signaling. The Sevenless (Sev) receptor tyrosine kinase hyper-activates MAP kinase in certain eye progenitor cells, leading to enhanced turnover of the Yan protein (Rebay and Rubin, 1995). This enables cells to differentiate into R7 photoreceptors (Voas and Rebay, 2004). When *sev* is mutated, cells completely fail to differentiate as R7 photoreceptors. This effect was readily apparent by staining for an R7-specific marker protein (Fig. 1F). However, slowing metabolism allowed a small but significant number of *sev* mutant cells to become R7 photoreceptors (Fig. 1G). Importantly, since the *sev* mutant makes no protein products (Banerjee et al., 1987), rescue of the mutant phenotype was not simply due to more functional Sev protein molecules being present in slowly metabolizing cells.

We also examined formation of other sensory organs for evidence of metabolic interactions. Large sensory bristles develop in a highly stereotypic pattern over the *Drosophila* body. The protein Senseless (Sens) transiently appears in a cluster of proneural cells before one cell is chosen to differentiate into a sensory bristle (Jafar-Nejad et al., 2003). MicroRNA miR-9a represses Sens protein expression, and *mir-9a* mutants frequently develop ectopic sensory bristles because this repression is missing (Fig. 2A,B) (Cassidy et al., 2013; Li et al., 2006). However, when *mir-9a* mutants had their IPCs ablated, errors in bristle number were greatly reduced (Fig. 2C).

**Figure 2.**
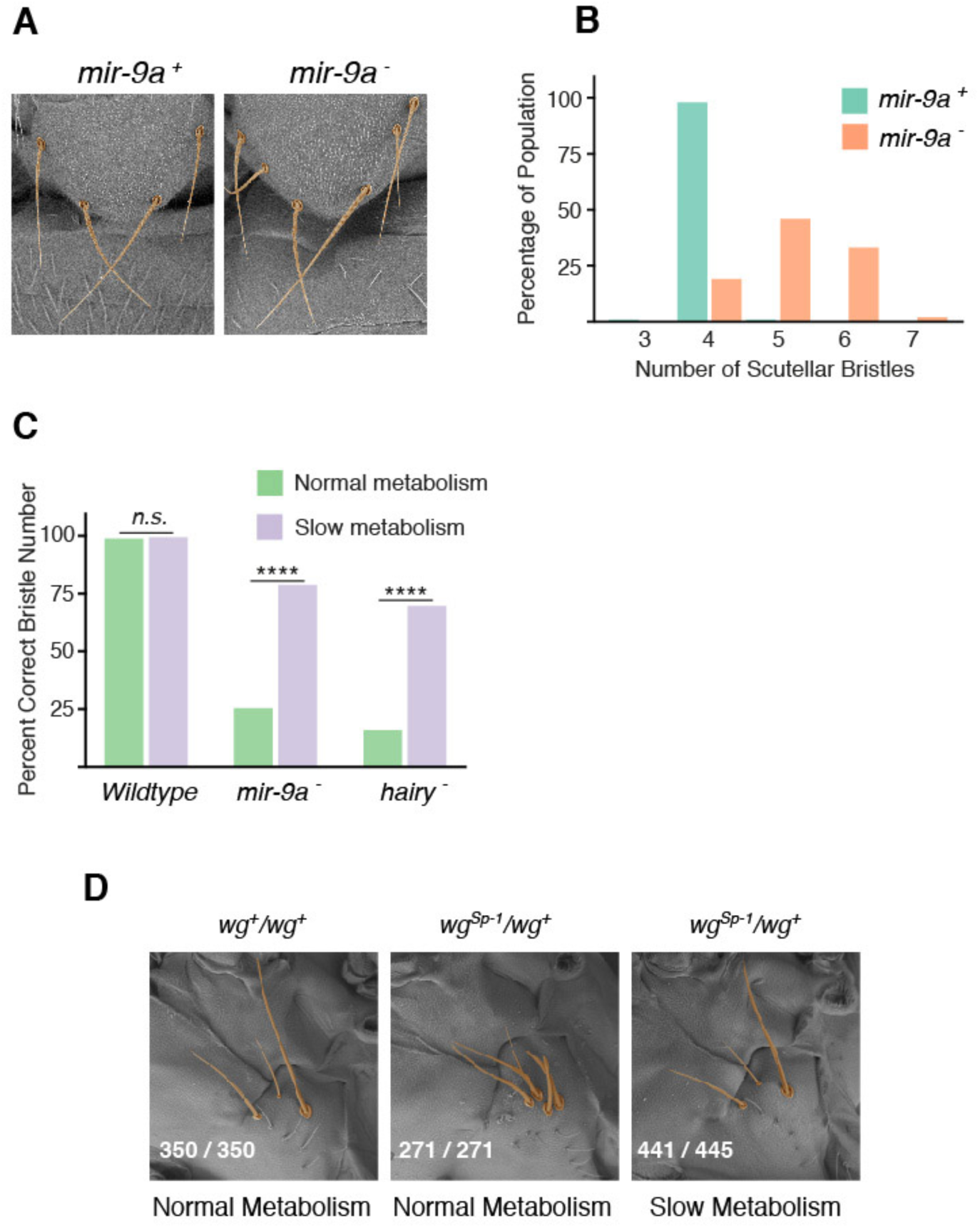
Sensory bristle developmental defects are rescued by slower energy metabolism. **(A)** Number of scutellar bristles is frequently greater than four in a *mir-9a* mutant whereas it is almost invariably four in wildtype. **(B)** Distribution of scutellar bristle numbers in wildtype and *mir-9a* mutant populations. Population sizes for wildtype and *mir-9a* were 301 and 222, respectively. The cumulative frequency distributions between wildtype and mutant were significantly different (p < 0.0001, Kolmogorov Smirnov test). **(C)** IPC ablation increases the proportion of *mir-9a* mutants and hairy mutants that have the wildtype number of scutellar bristles. ****, *p* < 0.0001; n.s., *p* > 0.05 **(D)** Under normal metabolic conditions, wg^sP-1^displays an increased number of sternopleural bristles. IPC ablation dramatically increases the number of mutant individuals with the wildtype number of three sternopleural bristles. Shown in each panel is the number of individuals with bristle number of three versus the total number of individuals scored. IPC ablation significantly suppresses the wg^sP-1^ mutant phenotype (*p* < 0.0001, Fishers exact test).

The protein Hairy directly represses transcription of the proneural genes *achaete* and *scute* during selection of cells for bristle fates (Van Doren et al., 1994). Mutation of *hairy* causes some individuals to develop ectopic large bristles. However, this effect of *hairy* mutation was strongly suppressed when energy metabolism was slowed (Fig. 2C). We saw a similar effect on a cis-regulatory module (CRM) that represses gene transcription. The *Sternopleural* (*Sp-1*) mutation is present in a CRM located on the 3’ side of the *wingless* (*wg*) gene (Neumann and Cohen, 1996), causing Wg misexpression and development of ectopic bristles (Neumann and Cohen, 1996). However, the ectopic bristle phenotype of the *wg^sp-1^* mutant was completely reversed under conditions of slowed energy metabolism (Fig. 2D).

### MicroRNAs Are Dispensable When Metabolism is Reduced

The mutations thus far examined affect diverse types of regulators, including microRNAs, transcription factors, and signaling molecules. Despite this diversity, all of the mutations have something in common: they affect repressive interactions between genes. To explore the prevalence of this relationship between gene repression and metabolism, we eliminated an entire family of regulatory repressors that control all stages of *Drosophila* development. The microRNA family is composed of 466 distinct microRNAs in *Drosophila melanogaster* (Kozomara and Griffiths-Jones, 2014). Virtually all microRNAs require Dicer-1 (Dcr-1) protein for their proper biosynthesis, and Ago1 protein as a partner to repress target gene expression (Carthew and Sontheimer, 2009). Protein-null mutations in either *dcr-1* or *ago1* genes are lethal (Pressman et al., 2012). We raised different null *dcr-1* mutants under conditions of slower energy metabolism, and found that many more animals survived development (Fig. 3A). *Ago1* null mutants are normally 100% embryonic lethal, but mutant lethality was broadly suppressed when animals slowly metabolize due to IPC ablation (Fig. 3B). The mutants survived to adulthood, and most survivors had normal eye and bristle patterns as well as other body structures, indicating the rescue of a massive array of developmental defects (Fig. 3C). Rescue could also be seen when Ago1 was specifically ablated in cells of the compound eye; eye development was strongly rescued by slower energy metabolism (Fig. 3D). Therefore, a major class of regulatory repressors is rendered non-essential when energy metabolism is slowed.

**Figure 3.**
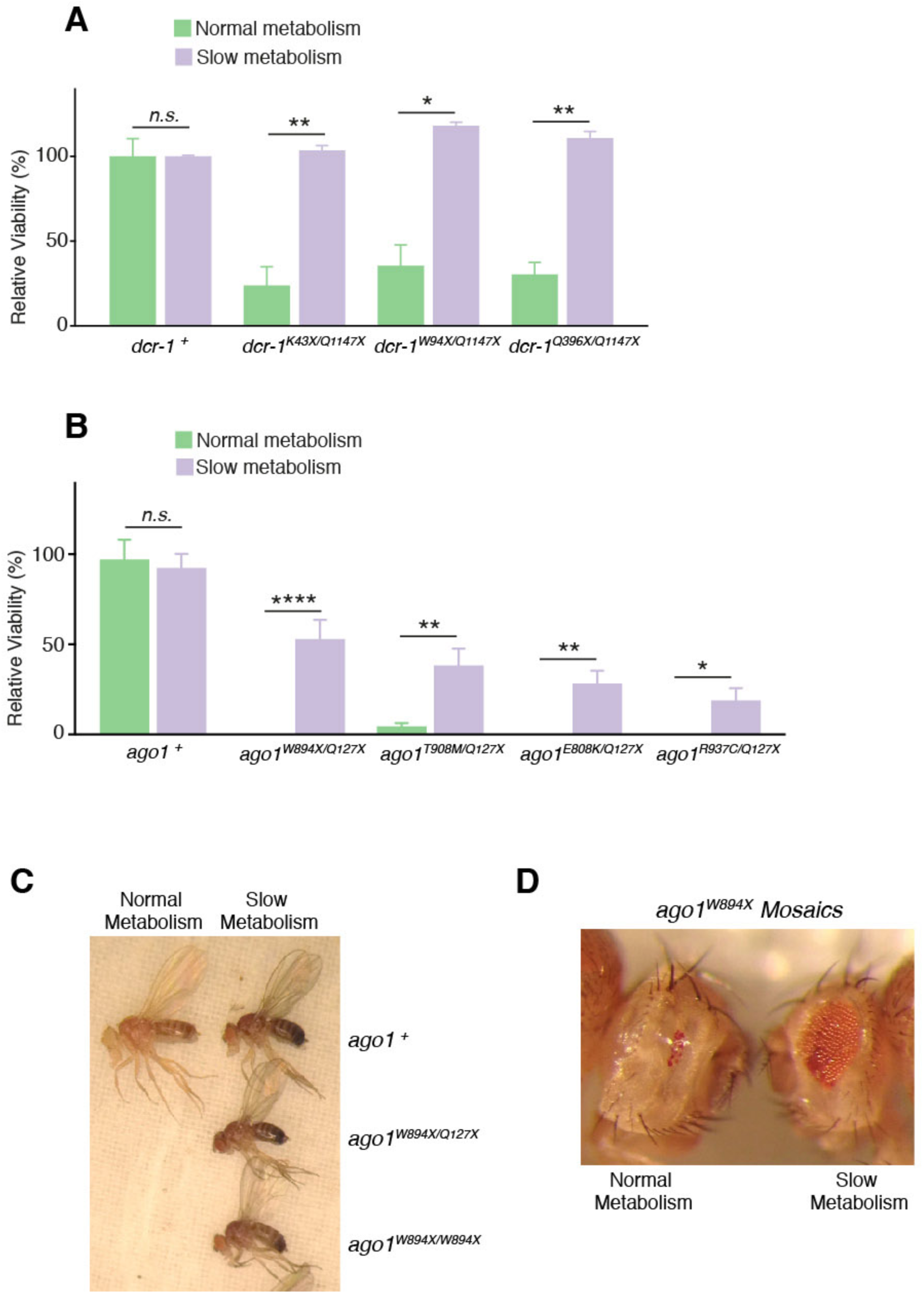
The microRNA family is dispensable when energy metabolism is slowed. **(A)** The pupal viability of various dcr-1 nonsense mutants is fully rescued when IPCs are ablated in the mutants. **(B)** Adult viability of various ago1 missense and nonsense mutants is rescued when IPCs are ablated in the mutants. **(C)** Representative ago1 adults with normal or slowed metabolism. **(D)** Genetically mosaic individuals with ago1+ bodies and *ago1^w894X^* mutant eyes. Left, representative individual with normal metabolism has almost no eye tissue (24/24 animals had this phenotype). Right, representative individual with slowed metabolism has rescued eye tissue. Of 70 such animals analyzed, 46 had this phenotype, 20 had normal eyes, and 4 had eyes that resembled the left animal. This is a statistically significant difference (*p*< 0.0001; Chi square with Yates’ correction). Error bars, s.d. ****, *p* < 0.0001; **, *p* < 0.01; *, *p* < 0.05; n.s., *p* > 0.05

### A Quantitative Model Describes the Relationship Between Metabolism and Developmental Error Frequency After Repressor Loss

We turned to computational modeling in order to elucidate the biochemical mechanism linking gene repression and metabolism. Because this relationship appears to apply to many GRNs operating at many stages of *Drosophila* development, we sought to model a general feature of dynamical systems rather than the specific regulatory mechanisms behind each of our experimental observations. We therefore explored the mechanism using a general modeling framework for GRNs premised on the progressive restriction of cell potential over time. Each step of restriction corresponds to a change in gene expression (Fig. 4A). A common property of such dynamics is that they are transient; gene products are synthesized, act, and then are eliminated until they are again needed in other cells. This property allows signaling molecules and transcription factors to be repeatedly used to build different body structures at different times.

**Figure 4.**
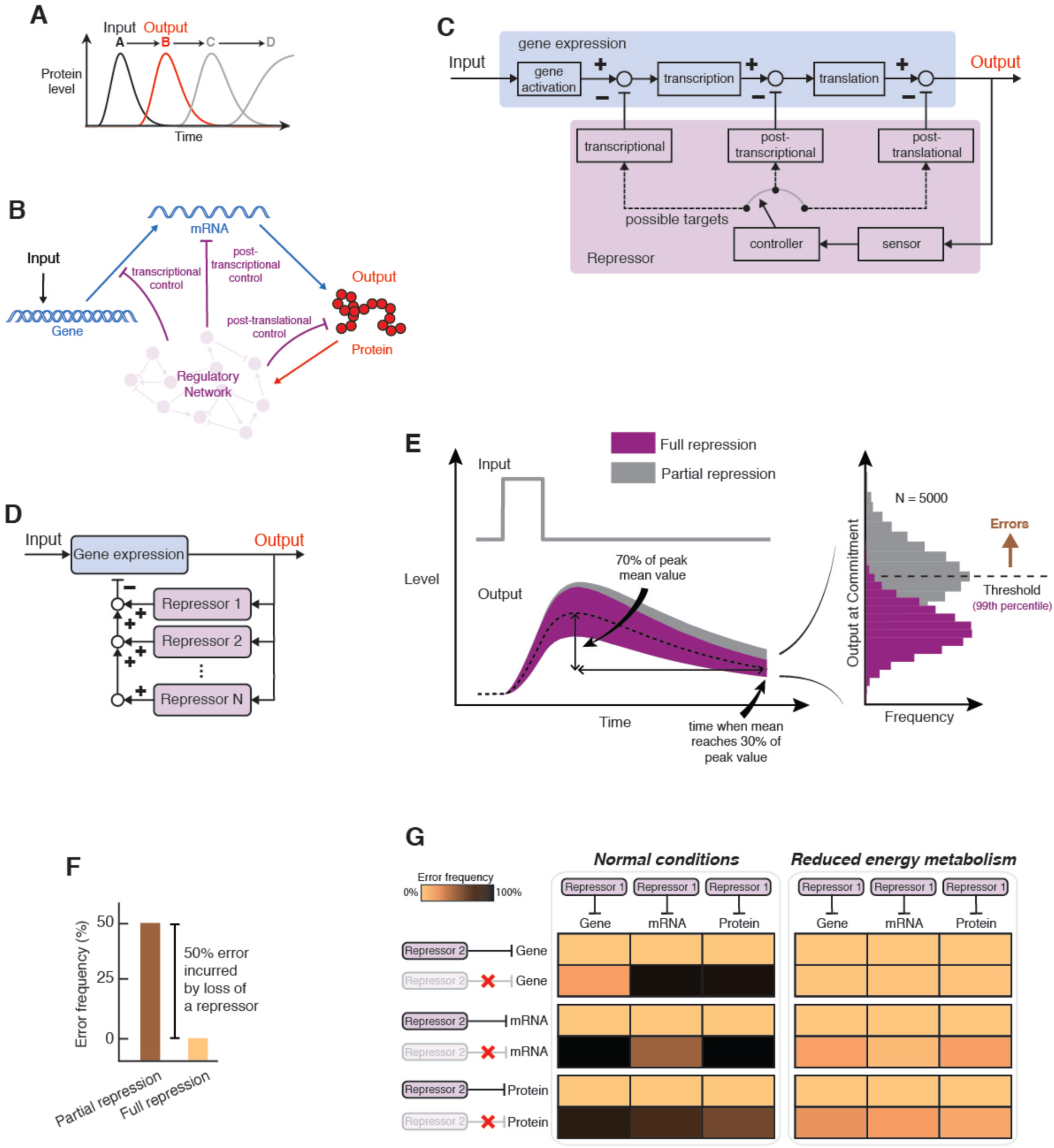
Modeling metabolism and gene regulation during cell fate determination. See also Figures S1 - S4. **(A)** A program of gene expression occurs as a single cell passes through a series of developmental states. The model focuses on transient expression of a single gene within a cascade of gene expression. A state change is defined as the induction of gene expression by upstream gene products (input) and the action of the gene product (output). **(B)** Schematic representation of the response to a transient input, which can be either an extracellular or intracellular signal. Gene expression output is subject to layers of negative regulation acting at the gene, transcript, and protein levels. **(C)** Control representation of a single feedback loop as depicted in B. Boxes contain transfer functions, open circles indicate summation points, and closed circles indicate exclusive switches for each repressor. **(D)** Protein expression may be subject to layers multiple repressors acting in parallel. **(E)** Simulation of protein levels in response to a transient input signal. One condition has two post-translational repressors in place (full repression) and the other condition has one repressor in place (partial repression). Shaded regions correspond to the 98% confidence band of simulated protein trajectories. We define the commitment time as the time needed for 99% of simulations with full repression to cross a predefined threshold. With partial repression, the protein levels take longer to decay, so fewer simulations cross the threshold within the defined commitment time. We interpret each failure of a simulated protein level to decay below the threshold in time as a developmental error. **(F)** Error frequency is greater with partial repression. **(G)** The model suggests that errors will occur more frequently under partial repression regardless of how repressors act on gene expression (left panel). However, partial repression imparts fewer developmental errors when ATP-dependent parameter values are reduced by 50% (right panel).

GRNs use layers of negative regulation to attenuate expression of target genes (Fig. 4B). When these targets are induced by exogenous stimuli, their timely attenuation ensures that protein expression remains transient. The resultant dynamics resemble a simple pulse response, and for this reason classical control theory provides a natural modeling framework (Figs. 4C and S1A). In our model, a system of cellular components monitors the relative abundance of a regulatory protein that dictates a cell fate transition. The particular components responsible for implementing control may remain unspecified. Our modeling framework therefore eschews molecular events in favor of minimizing complexity and preserving universality, but preserves the salient features of a detailed molecular mechanism and promotes quantitative predictions.

A stimulus (the input) activates synthesis of the regulatory protein (the output). Acting in parallel, one or more feedback control elements sense the increase in protein levels and act to down-regulate it, either at the level of gene transcription, mRNA stability, or protein stability. These control elements can be thought of as independent repressors working in parallel to bring the protein level back to a basal steady-state (Fig. 4D). Protein expression follows a biphasic trajectory after reception of a transient stimulus (Fig. 4E, left panel). If there were no noise or variability, the protein level would be deterministic over time. However, protein dynamics vary because gene expression is noisy (Arias and Hayward, 2006), something that can be captured in the model simulations by incorporating intrinsic noise.

We then devised a scheme to relate protein expression dynamics to the likelihood of a successful developmental outcome. We define success as the ability of a GRN to attenuate protein expression in a timely manner, thus keeping pace with parallel components of the developmental program by triggering subsequent developmental events. We quantified errors in developmental outcome by defining a threshold that the output protein level must cross before a subsequent event can be triggered. Protein levels exceeding the threshold constitute errors in developmental outcome (Fig. 4E, right panel). Notably, such errors become more frequent when one repressor is removed (Fig. 4F). This property is observed over a broad range of parameter values, regardless of the manner in which repressors act, or the value at which the threshold is established (Fig. S1B-C).

The modeling framework allowed us to ask whether multiple layers of repression are less important for developmental outcome when energy metabolism is reduced. To answer this question, we halved the rate parameters of each ATP-utilizing reaction to reflect conditions of reduced energy metabolism. Although ATP content remains fairly constant in cells facing limited respiration, the fluxes of ATP synthesis and turnover are affected, manifesting in altered ratios of ATP to ADP and free phosphate (Brown, 1992). Anabolic processes such as protein synthesis are highly dependent on the ATP/ADP ratio (Atkinson, 1977). When we halved ATP-dependent rate parameters and compared model results from full versus partial repression, we observed that error frequency in developmental outcome did not increase when a repressor was lost (Fig. 4G). This insensitivity to loss of a repressor persisted whether repression was transcriptional, post-transcriptional, or post-translational. The effect was observed across a wide range of model parameter values (Figs. S2A,B), irrespective of where the threshold was set (Fig. S2C), and regardless of whether a basal stimulus was present (Fig. S2D). In many cases the effect remained modestly apparent when the stimulus duration was extended to maintain comparable protein levels under conditions of reduced energy metabolism (Fig. S2E). In general, our modeling framework suggests that the frequency of developmental errors is less sensitive to changes in repression when energy metabolism is reduced.

Our modeling framework for regulated gene expression promotes simplicity at the expense of two notable limitations. First, the number of transcriptionally active sites within a cell is limited by gene copy number, but the activated-DNA state in our initial linear model was unbounded. To test whether error frequency suppression persists when an upper bound on gene activity is introduced, we considered a simple two-state transcription model. Despite the limitation placed on gene activity, error frequencies remained broadly suppressed when ATP-dependent rate parameters were reduced (Fig. S2F).

Second, gene expression models frequently utilize cooperative kinetics in order to reproduce the nonlinearities and thresholds encountered in transcriptional regulation. We captured these dynamics by reformulating our gene expression modeling framework in terms of Hill kinetics. A parameter sweep revealed that despite the incorporation of cooperative kinetics, error frequencies remained elevated under normal conditions and broadly suppressed when ATP-dependent rate parameters are reduced (Fig. S2G).

Reduced glucose consumption by cells might not only limit ATP fluxes, but also hinder the synthesis of nucleotide and amino acid precursors required for RNA and protein synthesis. To simulate this scenario, we specifically reduced the rate parameters for RNA and protein production. Constraining these synthesis rates also suppressed the rise in error frequency when a repressor was lost (Fig. S2H).

Overall, our modeling framework is fully consistent with our experimental observations that multiple layers of repression cease to be important for developmental success under conditions of reduced carbon and energy metabolism. Furthermore, the modeling framework suggests that the phenotype suppression phenomenon may be driven by differences in protein expression dynamics that are dependent on metabolic conditions.

### Protein Expression Dynamics After Partial Repressor Loss

We quantified the extent to which the expression dynamics underlying developmental events were affected in our model. We first constructed a 98% confidence band around the set of trajectories simulated with the full complement of repressors (Fig. S3A). This confidence band provides lower and upper bounds for the expected protein level. We then evaluated the fraction of trajectories simulated with one repressor missing that fell above or below the confidence band at each point in time. Averaging these values across the time course yields a single metric that reflects the extent to which protein dynamics are affected by the loss of a repressor (Fig. S3B). We evaluated this metric for a scenario in which an auxiliary post-transcriptional repressor, akin to a microRNA, is lost. Using typical metabolic parameters, 78% of trajectories simulated without the post-transcriptional repressor exceed the confidence band generated under full repression (Fig. 5A). This overexpression effect is highly robust to parameter variation in the model (Fig. S3C). When ATP-dependent parameters were halved, only 16% of trajectories exceeded the confidence band (Fig. 5B). The strong diminishment of overexpression under low metabolic conditions was robust to extensive parameter variation (Fig. S3D). These results led us to predict that protein expression dynamics would be much less sensitive to repressor loss if we reduced metabolic rate.

**Figure 5.**
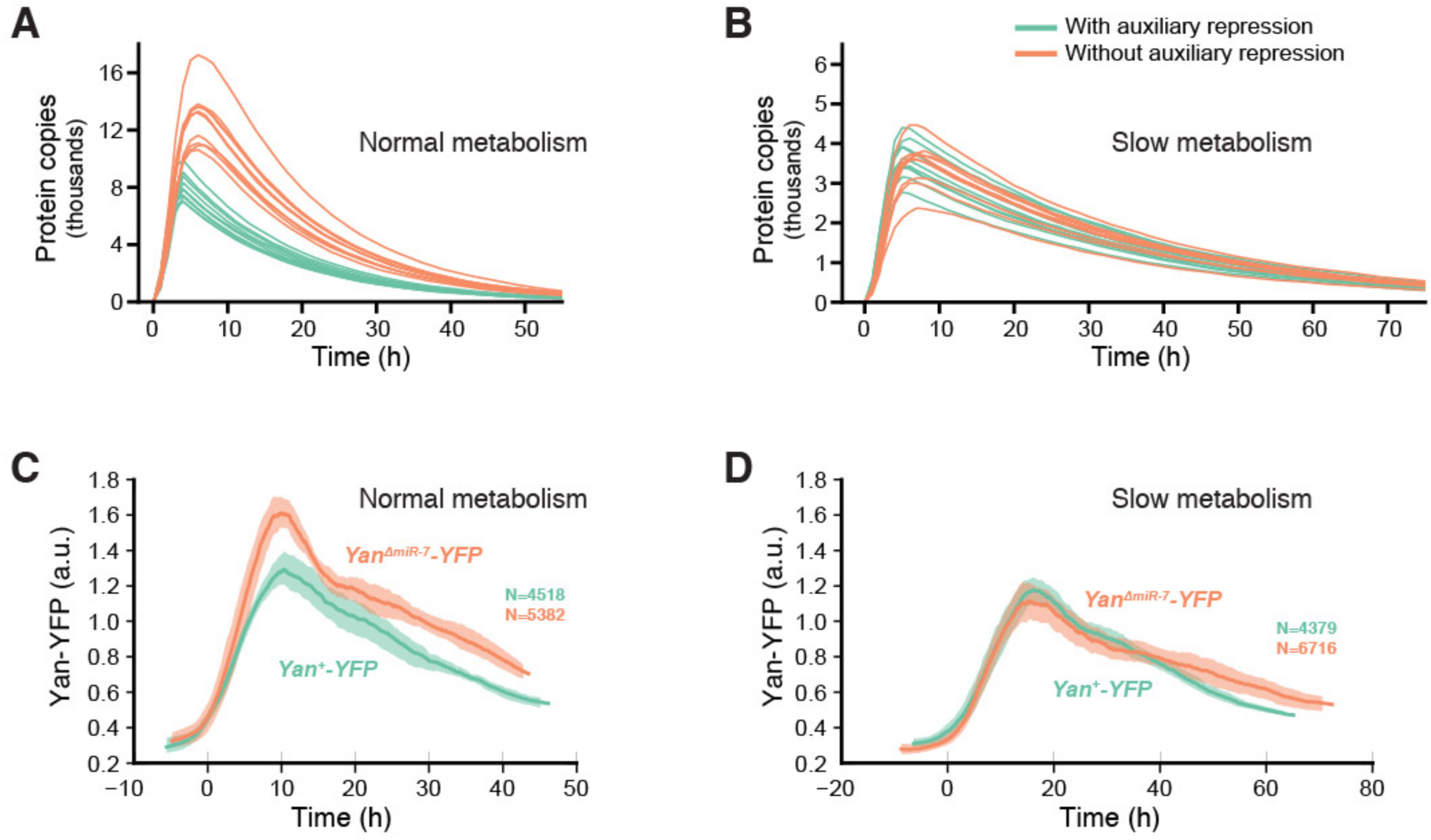
Expressiori dynamics are resistant to repressor loss when energy metabolism is reduced. **(A,B)** Simulated expression of target protein output when it is under control of an auxiliary post-transcriptional repressor (green) or not under control of the repressor (orange). All simulations (green and orange) are also under control of a constitutive repressor. Shown are ten randomly-chosen samples from a total population of 5000 trajectories for each condition. **(A)** Simulations performed with normal ATP-dependent reaction rates. **(B)** Simulations performed following a 50% reduction in the rate of ATP-dependent reactions. **(C,D)** Yan-YFP protein dynamics in eye disc progenitor cells. Time 0 marks the time at which Yan-YFP induction occurs. Solid lines are moving line averages. Shaded regions denote 95% confidence intervals. Each line average is calculated from a composite of measurements of between 4,379 and 6,716 cells. **(C)** Yan-YFP dynamics for wildtype *Yan-YFP* and mutant *Yar^ΔmiR-7^-YFP* genes under normal metabolic conditions. **(D)** Yan-YFP dynamics for wildtype and mutant genes when the I PCs have been ablated.

We experimentally tested these predictions by measuring the expression dynamics of a key developmental regulatory protein, Yan. Yan exhibits pulsatile dynamics in the larval eye disc, where its expression is induced by a morphogenetic furrow that traverses the eye disc. Eye disc cells located in the morphogenetic furrow rapidly upregulate Yan protein abundance, as quantified by a YFP-tagged version (Pelaez et al., 2015). Yan levels then gradually decay back to initial conditions within these cells, thus exhibiting pulsatile dynamics. We compared Yan-YFP dynamics in eye disc cells from normally metabolizing larvae and larvae with ablated IPCs (Fig. 5C,D). The same pulsatile dynamics were observed in both, but the amplitude of the pulse was slightly reduced and the duration was extended when metabolism was slower. Similar trends were predicted in silico (Fig. 5A,B).

In the eye disc, Yan expression is repressed by the microRNA miR-7 (Li and Carthew, 2005). There are four binding sites for miR-7 in the 3'UTR of *yan* mRNA, and their mutation causes de-repression of Yan output. We eliminated miR-7 repression of *Yan-YFP* by mutating the four binding sites in the 3'UTR of *Yan-YFP* mRNA to make *Yan^ΔmiR-7^-YFP*. In normally metabolizing eye discs, Yan-YFP protein made from the mutated gene pulsed with greater amplitude and showed impaired decay when compared to Yan-YFP made from the wildtype gene (Fig. 5C). These dynamics recapitulate the effect of repressor loss predicted by our model (Fig. 5A). In contrast, Yan-YFP made from the mutated gene showed similar dynamics to protein made from the wildtype gene when metabolism was slowed (Fig. 5D). This behavior clearly resembled the simulated dynamics under conditions of reduced energy metabolism (Fig. 5B).

These measurements demonstrate that miR-7 has little to no impact on Yan expression dynamics when metabolism is slowed, and are consistent with the observed suppression of developmental errors when the same repressor is lost in the eye. The breadth of our model predictions further suggests that these effects are generalizable to other genes and repressors.

### Effect of Full Repression Loss

Our modeling framework is consistent with the hypothesis that multiple weak repressors allow GRN dynamics to faithfully couple to variable energy metabolism, with fewer repressors required when metabolic conditions are reduced. We then asked whether repression is needed at all under such conditions. We studied a model with a full complement of negative control elements and compared the results to a scenario in which all control elements were removed (Fig. 6A). Error frequencies approached 100% under normal growth conditions. While expression dynamics were visibly less affected by repressor loss when ATP-dependent parameters were reduced, the error frequency remained very high (Fig. 6B). These results suggest that there are limits to the severity of perturbations for which reductions in energy metabolism can compensate, and reducing energy metabolism does not eliminate the need for gene repression altogether.

**Figure 6.**
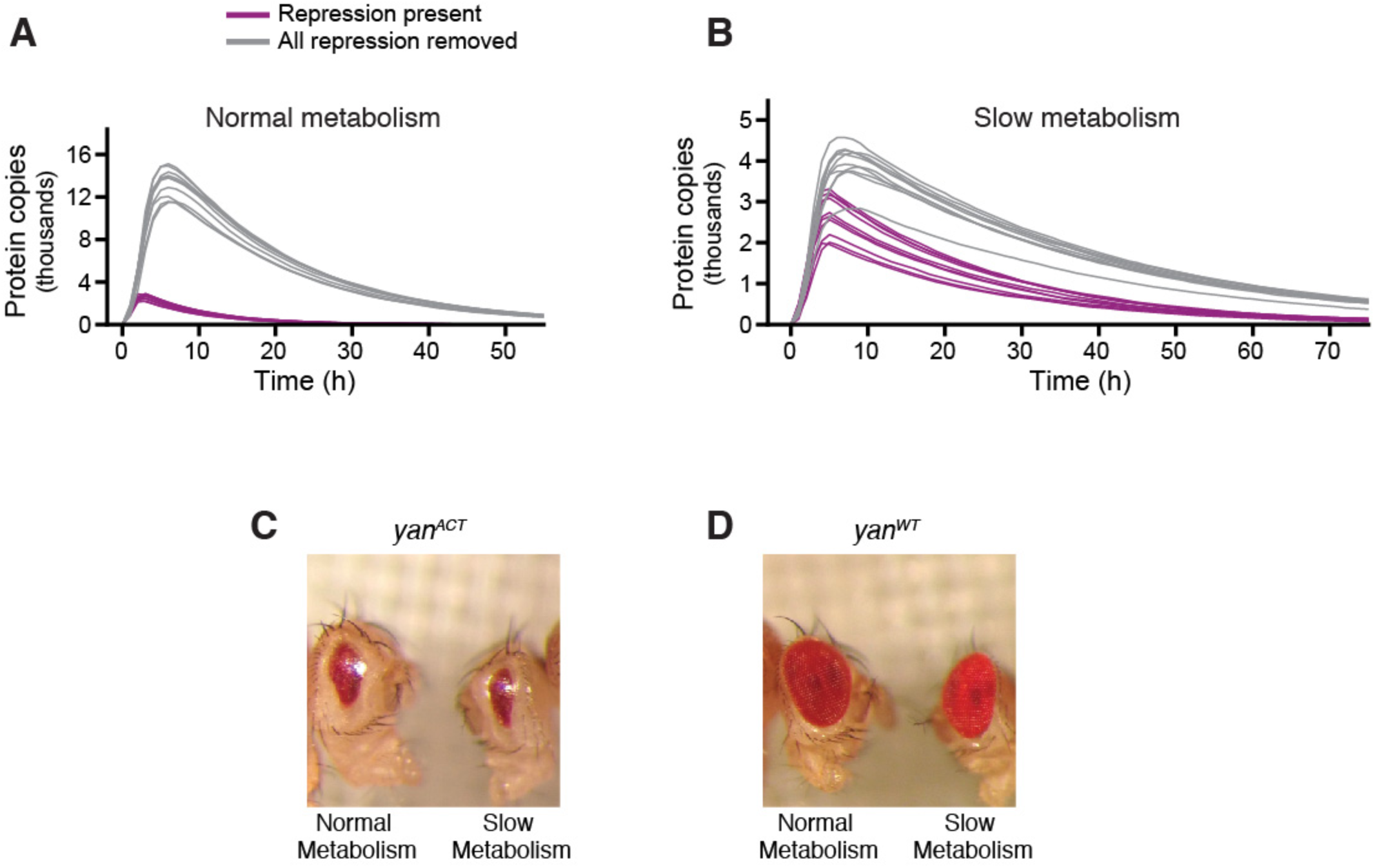
Reduced energy metabolism cannot compensate for complete loss of repression. **(A,B)** Simulated expression of protein output both with (purple) and without (grey) any repression of the target gene. Shown are ten randomly chosen samples from a total population of 5000 trajectories for each condition. Error frequencies exceed 99% irrespective of metabolic conditions. **(A)** Simulations performed under normal conditions. **(B)** Simulations performed following a 50% reduction in the rate of ATP-dependent reactions. **(C)** Loss of eye tissue in a *yan^ACT^* mutant is not suppressed by slower metabolism. Representative individuals were taken from N > 100 individuals for each condition. **(D)** Eye patterning in a *yan^WT^* control is not affected by slower metabolism. Representative individuals were taken from N > 100 individuals for each condition.

To test this prediction, we expressed in the eye a *yan* mutant transgene that is insensitive to all known repression of *yan* transcription, mRNA stability and protein stability (Rebay and Rubin, 1995). The *Yan^Act^* mutant adults had severely disrupted compound eye patterning (Fig. 6C). This mutant eye phenotype was not suppressed by ablation of the animals' IPC cells. Wildtype *yan* transgenic adults with normal eye patterning were also unaffected by IPC ablation (Fig. 6D).

### Limiting Protein Synthesis Reduces the Need for Repressors

The coupling of developmental dynamics to time can be explored with other aspects of metabolism. In particular, protein synthesis is an important determinant of rates of growth and development (Lempiainen and Shore, 2009). We used our modeling framework to investigate the impact of a twofold reduction in overall protein synthesis rate on GRN dynamics. The model suggests that expression dynamics are less affected and fewer developmental errors are incurred by loss of a repressor when protein synthesis rates are reduced (Fig. 7A and Figs. S4A-G).

**Figure 7.**
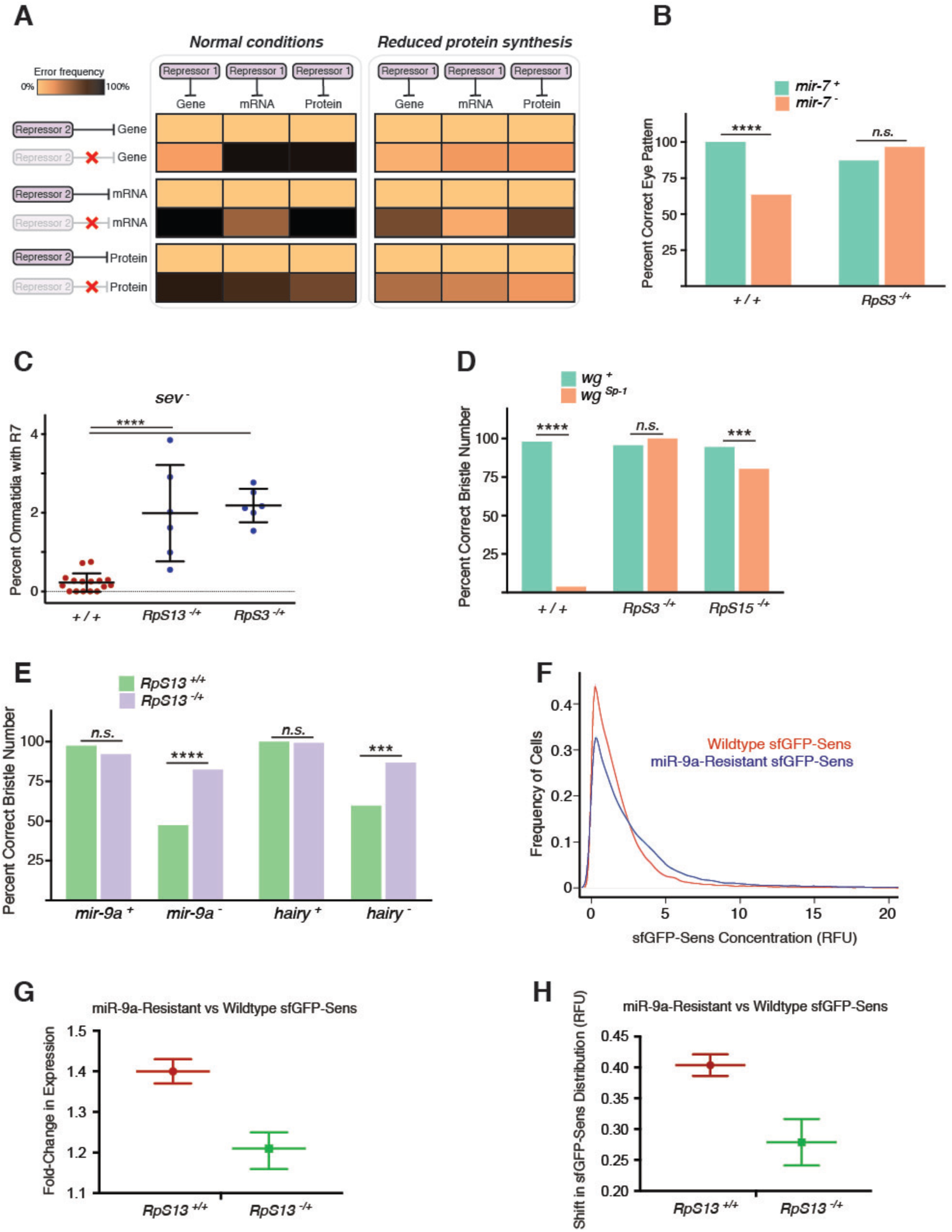
Reducing ribosome number rescues loss of repressors on sensory organ development and expression dynamics. **(A)** The model predicts increased frequency of error with partial repression regardless of how auxiliary repressors act on gene expression (left panel). However, partial repression induces fewer errors when protein synthesis-dependent parameter values are reduced by 50% (right panel). **(B)** Loss of *mir-7* does not cause adult eye mispatterning when *RpS3* is heterozygous mutant. **(C)** *sev* mutants have more R7-positive ommatidia when either *RpS3* or *RpS13* are heterozygous mutant. Each datapoint represents one eye sample, and between 481 and 837 ommatidia were scored for R7 cells within each eye sample. **(D)** wg^sP-1^heterozygous individuals that are also heterozygous mutant for different RpS genes have sternopleural bristle numbers more similar to wildtype. **(E)** Developmental accuracy is recovered for both mir-9a and hairy mutants that are also heterozygous mutant for *RpS13*. For all panels in B-E, error bars, s.d. ****, *p* > 0.0001; ***, *p* < 0.001; n.s., *p* > 0.05. **(F)** Frequency distribution of sfGFP-Sens protein level in cells bordering the wing margin of white prepupal wing discs. Shown are distributions of cells expressing either wildtype *sfGFP*-sens or *sfGFP*-sens in which miR-9a binding sites have been mutated. Shaded regions denote 95% confidence intervals. Each line is calculated from a composite of measurements of > 18,000 cells. **(G)** The fold-change in median *sfGFP-sens* expression caused by miR-9a binding site mutations. Measurements were taken in *RpS13* wildtype and heterozygous mutant backgrounds. Shown are 99% confidence intervals for the estimated fold-change. **(H)** The shift in the fluorescence distribution of sfGFP-Sens-positive cells caused by miR-9a binding site mutations, as determined from a Mann-Whitney-Wilcoxon test. The upward shift is smaller in an *RpS13* heterozygous mutant background. Shown are 95% confidence intervals for the shift.

We tested this model prediction by genetically reducing the abundance of cytoribosomes in all cells in *Drosophila*. We made use of loss-of-function mutations in genes encoding various ribosomal proteins (RPs), which cause the “Minute” syndrome of dominant, haploinsufficient phenotypes, including slower growth and development (Marygold et al., 2007; Sæbøe-larssen et al., 1998). Heterozygous *RP* mutants reduce the number of ribosomes per cell by approximately 50%, and a total of 64 *RP* genes exhibit a Minute syndrome when mutated. We selected a subset of these genes to reduce ribosome number. We combined heterozygous *RP* mutants with the repressor mutations we had previously studied. In all cases, the *RP* mutants suppressed the developmental phenotypes of mutations in *wg*, *mir-7*, *sev, hairy*, and *mir-9a* (Fig. 7B-E). This error frequency suppression was precisely the result predicted by our modeling.

We also tested whether expression dynamics are affected by repressor loss under limiting translation conditions. The Sens protein is transiently expressed in proneural cells during selection of sensory bristle fates in the imaginal wing disc (Nolo et al., 2000). Bordering the presumptive wing margin, stripes of proneural cells express Sens protein over a spectrum of levels, reflecting heterogeneity in Wg and Notch regulation of its expression (Jafar-Nejad et al., 2006; Quan and Hassan, 2005). Moreover, miR-9a weakly represses *sens* expression in these cells (Li et al., 2006). We recombineered a 19 kb *sens* transgene, amino-terminally tagged with superfold GFP (sfGFP), that functionally replaced the endogenous *sens* gene (Cassidy et al., 2013; Venken et al., 2006). Quantitative measurement of sfGFP fluorescence in individual proneural cells yielded the expected distribution of *sens* expression (Fig. 7F). We compared this distribution to one derived from individuals expressing a mutated sfGFP-*sens* transgene in which its miR-9a binding sites had been mutated (Cassidy et al., 2013). Mutation of the miR-9a binding sites in *sfGFP-sens* shifted the fluorescence distribution (Fig. 7F), and resulted in an average 1.4-fold increase in sfGFP-Sens levels (Fig. 7G). We then tested the effects of miR-9a on *sfGFP-sens* expression when the *RpS13* gene was heterozygous mutant. Strikingly, loss of miR-9a regulation had less effect on sfGFP-Sens protein levels when ribosome numbers were reduced (Fig. 7G,H). This behavior clearly resembled the simulated dynamics under conditions of reduced protein synthesis (Fig. S4B).

## DISCUSSION

Growth and development are fuelled by metabolism. This means that the tempo of development depends on metabolic rate. Thus, the dynamics of developmental gene expression must faithfully adjust to a variable time scale. We have shown that multi-layered weak repression within GRNs plays an unexpected function in synchronizing gene expression dynamics with the variable pace of the developmental program. Multiple repressors are required for accelerated development when metabolism is high, and they become functionally redundant when metabolism is low. Multiple repressors therefore allow for reliable development across a broader range of metabolic conditions than would otherwise be tolerated.

Our model explains long-standing observations linking nutrient limitation to suppression of mutant phenotypes (Morgan, 1915; Morgan, 1929). Presumably, such mutations cripple regulatory genes acting on developmental GRNs. Our model might also offer an explanation as to why animals that undergo above-normal growth exhibit compromised development (Arendt, 1997; Metcalfe and Monaghan, 2001). Wildtype GRNs might function across a limited range of metabolism, with functionality breaking down when metabolism exceeds that range.

Our varied analyses suggest that this relationship between metabolism and repression is ubiquitous. We found that the entire family of 466 microRNAs in *Drosophila melanogaster* can become functionally dispensable when energy metabolism is slowed. The extensive literature on microRNA function in *Drosophila* implicates them in practically all facets of the fruit fly’s life (Bushati and Cohen, 2007; Carthew et al., 2017). Various explanations have been provided for why this family of weak repressors has flourished in the animal kingdom, chief among them the idea that they act as buffers for gene expression (Ebert and Sharp, 2012). We now posit that microRNAs also provide broad and flexible coupling of many developmental processes to variable timescales resulting from fluctuations in metabolism.

There is an alternative mechanism to explain phenotype suppression by reduced metabolism. This mechanism relies on a steady-state and not dynamical perspective of gene expression. Genome-wide gene expression patterns could conceivably change with organismal growth rate. This is the case for chemostat-grown yeast cells, where the expression of 27% of all genes correlates with growth rate (Brauer et al., 2008). Most genes associated with stress response are overexpressed when cells grow at a slow rate (Brauer et al., 2008; Lu et al., 2009). Such differential gene expression could globally modulate dynamical processes such as protein folding and turnover, among others, and thereby attenuate phenotypes of genetic mutations. Abundance of molecular chaperones has been found to affect the penetrance of diverse gene mutations in *C. elegans* and *Drosophila* (Casanueva et al., 2012; Rutherford and Lindquist, 1998). However, these global effects do not explain why gene expression dynamics are conditionally dependent upon mutations in regulatory genes. We found that repression of Yan and Sens dynamics by microRNAs become more redundant when metabolic rates are slowed.

Metabolic rate increases exponentially with temperature as described by the Arrhenius equation (Zuo et al., 2011), resulting in an indirect temperature dependence of developmental tempo (Gillooly et al., 2002). Temperature also directly affects the rates of reactions within developmental GRNs (Zuo et al., 2011), yet developmental outcomes are generally robust to fluctuations in temperature across a limited range. Various molecular mechanisms have been invoked to explain this robustness. These include chaperones that create large protein-folding reservoirs (Jarosz and Lindquist, 2010; Rutherford and Lindquist, 1998), and regulatory circuits within interaction networks (Li et al., 2009). Our model suggests a complementary mechanism for developmental robustness against temperature variation. By coupling gene expression dynamics with metabolism, weak repressors might neutralize the metabolic effects of temperature on developmental tempo. Indeed, loss of miR-9a regulation is less impactful on sensory organ development if the growth temperature is lowered (Cassidy et al., 2013). Likewise, raising animals under lowered temperatures can suppress the phenotypes of mutations that are not classical *ts* alleles (Child, 1935; Krafka, 1920; Lewis et al., 1980; Villee, 1943).

Metabolic conditions drive variation about the intrinsic developmental tempo of each species. We have shown that layered weak repression within GRNs enables these fluctuations to occur without causing developmental errors. Metabolic conditions change in both space and time. Perhaps the selective advantage of a reliable developmental outcome amidst variable environmental conditions is a driving force in the evolution of gene regulatory networks.

## ACKNOWLEDGEMENTS

We thank Jon Braverman for discussions and insights, the Bloomington Drosophila Stock Center and the Developmental Studies Hybridoma Bank for reagents. We also thank Alec Victorsen and Jennifer Moran for help in Yan transgene recombineering, and Diana Posadas and Hemanth Potluri for help in Sens transgene recombineering. Jessica Hornick and the Biological Imaging Facility are acknowledged. We acknowledge support from the Chicago Biomedical Consortium (J.J.C., N.P.), the Malkin and Rappaport Foundations (J.J.C.), the Northwestern Data Science Initiative (R.G.), a John and Leslie McQuown Gift (L.A.N.A.), the NSF (1764421, L.A.N.A., N.B. and R.W.C), the Simons Foundation (597491 L.A.N.A., N.B. and R.W.C), and the NIH (T32 GM008061, J.J.C., B.E.), (T32 CA080621, R.B.), (P50 GM81892 and R35 GM118144, R.W.C).

## AUTHOR CONTRIBUTIONS

The experimental work was conceived by J.J.C. and R.W.C. The genetic experiments classifying phenotypes were performed by J.J.C., B.E., and A.B., under the supervision of R.W.C. N.P. designed and built the *Yan-YFP^ΔmiR-7^* mutant line, and R.B. performed the imaging and analysis of Yan-YFP expression, with assistance from S.B, and under the guidance of R.W.C. R.G. built the *sfGFP-sens* wildtype and mutant stocks with *RpS13*, and she performed all imaging and analysis of sfGFP-Sens expression. The modeling was conceived by S.B., N.B., and L.A.N.A. All model analysis was performed by S.B. under the supervision of N.B. and L.A.N.A. The manuscript was written by S.B. and R.W.C. with input from all authors.

## COMPETING INTERESTS

The authors declare no competing financial interests.

## CORRESPONDING AUTHORS

Correspondence to Richard Carthew and Luis Amaral.

## STAR METHODS

### EXPERIMENTAL MODEL AND SUBJECT DETAILS

For all experiments, *Drosophila melanogaster* was raised using standard lab conditions and food. Stocks were either obtained from the Bloomington Stock Center, from listed labs, or were derived in our laboratory (RWC). A list of all mutants and transgenics used in this study is in the Key Resources Table.

Experiments were performed using either homozygous mutant animals or trans-heterozygous mutants. Trans-heterozygous allele combinations used were:

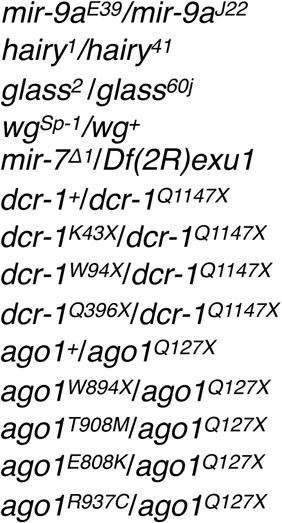

To genetically ablate the insulin producing cells (IPCs) of the brain, *yw* animals were constructed bearing an *ILP2-GAL4* gene on chromosome III and a *UAS-Reaper* (*Rpr*) gene on chromosome I or II. *Rpr* is a pro-apoptotic gene that is sufficient to kill cells in which it is expressed (Lohmann et al., 2002). *ILP2-GAL4* fuses the *insulin-like peptide 2* gene promoter to GAL4, and specifically drives its expression in brain IPCs (Rulifson et al., 2002). Examination of *ILP2-GAL4 UAS-Rpr* larval brains showed that they almost completely lacked IPCs (data not shown). Previous studies found that IPC-deficient adults are normally proportioned but of smaller size (Rulifson et al., 2002). It takes almost twice the length of time to complete juvenile development, and juveniles have a 40% elevation in blood glucose, consistent with these insulin-like peptides being essential regulators of energy metabolism in *Drosophila* (Rulifson et al., 2002). We confirmed that this method of IPC ablation results in small but normally proportioned adults, and it takes almost twice the normal time to develop into adults (Fig. 1B,C). For all wildtype controls, we tested animals bearing either the *ILP2-GAL4* or *UAS-Rpr* gene in their genomes.

To reduce levels of cytoribosomes in cells, we made use of loss-of-function mutations in genes encoding various ribosomal proteins (RPs), which cause the “Minute” syndrome of dominant, haploinsufficient phenotypes, including prolonged development (Sæbøe-larssen et al., 1998). A total of 64 *RP* genes exhibit a Minute syndrome when mutated (Marygold et al., 2007). We selected a subset of these genes to reduce ribosomes. Since one of these, *RpS3*, encodes an RP that also functions in DNA repair (Graifer et al., 2014), we tested it along with other *RP* genes in certain genetic experiments. The mutations used were: *RpS3^Plac92^* (Sæbøe-larssen et al., 1998), *RpS3^2^* (Ferrus, 1975), *RpS13^1^* (Sæbøe-larssen et al., 1998), and *RpS15^M(2)53^* (Golic and Golic, 1996). For wildtype controls, animals were *w^1118^*.

The recombineered *Yan-YFP* BAC transgene was previously described (Webber et al., 2013). We modified the gene by site-directed recombineering to mutate the four identified miR-7 binding sites within the *yan* (*aop*) gene (Li and Carthew, 2005). The binding sites and the mutations are listed below. The seed sequence is highlighted in red. The sequence of the mutations, which are localized to the seeds, are shown in green.

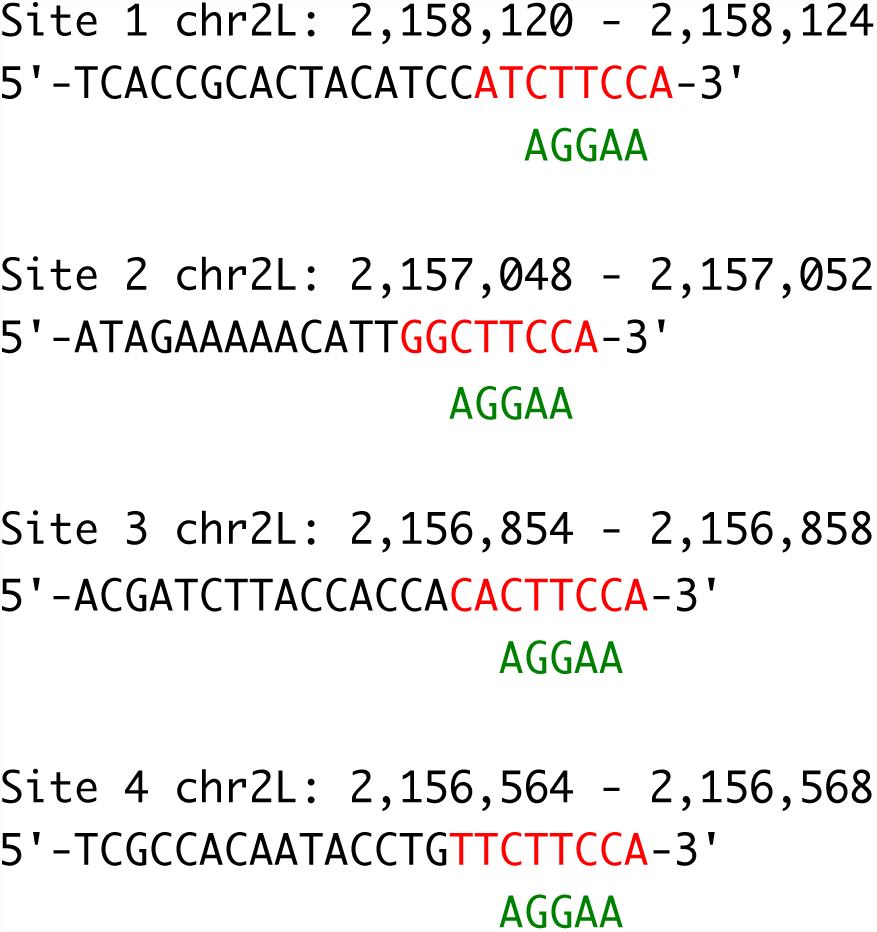

The mutated transgene (*Yan^ΔmiR-7^-YFP*) was shuttled into the P[acman] vector (Venken et al., 2006), and inserted into the same genomic landing site on chromosome 3 (attP2) as *Yan-YFP*. One copy of the *His2Av-mRFP* transgene was recombined with the *Yan^ΔmiR-7^-YFP* or *Yan-YFP* transgene in order to normalize YFP expression to a housekeeping protein, in this case histone H2A (Pelaez et al., 2015). The *His2Av-mRFP Yan-YFP* (*Yan^ΔmiR-7^-YFP*) chromosome was homozygosed, and placed in a *yan^ER443^ / yan^E884^* mutant background so that the endogenous *yan* gene did not make any protein.

The recombineered *sfGFP-sens* BAC transgene was generated as described (Cassidy et al., 2013), and the transgene was landed in the genome at VK37 (22A3). The transgene was mutated by site-directed recombineering as described (Cassidy et al., 2013) to delete the two miR-9a binding sites within the *sens* gene (*sfGFP-sens^m1m2^*). This transgene was also landed at VK37. The *sfGFP-sens* (*sfGFP-sens^m1m2^*) chromosome was homozygosed, and placed in a *sensE1* null mutant background to ensure that endogenous *sens* did not make any protein.

All experiments used female animals unless stated otherwise.

## METHOD DETAILS

### R7 Cell Analysis in the Eye

Individuals were synchronized at the larval-pupal transition, and incubated for a further 48 hours at 23°C. Eyes were dissected from pupae, and were fixed for 40 min in 4% paraformaldehyde/PBS. They were permeabilized by incubation in PBS + 0.1% Triton-X100 (PBST) and co-incubated with mouse anti-Prospero (1:10 in PBST, MR1A MAb, Developmental Studies Hybridoma Bank) to stain R7 and bristle cells plus rat anti-Elav (1:10 in PBST, 7E8A10 MAb, Developmental Studies Hybridoma Bank) to stain all R cells. After 60 min, eyes were washed 3 times in PBST and incubated for 60 min in goat anti-mouse Alexa546 and goat anti-rat Alexa633 (1:100 in PBST, Invitrogen). Eyes were washed 3 times in PBST, cleared in Vectashield (Vector Labs), and mounted for microscopy. Samples were scanned and imaged in a Leica SP5 confocal microscopy system. *Drosophila* compound eyes have approximately 800 ommatidia. We scored all ommatidia for each imaged eye sample. The number of scored ommatidia per sample ranged between 481 and 837 (with a median of 594). Fewer than 800 ommatidia were scored per sample because in most cases, some eye tissue was lost during dissection and handling.

### Relative Viability

Females bearing either a *dcr-1^Q1147X^* or *ago1^Q127X^* mutant chromosome over a balancer chromosome were crossed to males bearing mutant *dcr-1* or *ago1* chromosomes over a balancer chromosome. F1 progeny were raised and the numbers of animals that reached either pupal or adult stage were tallied. If the non-balancer chromosome is 100% viable when homozygous, then 33.33% of the F1 progeny would not carry a balancer chromosome. We calculated viability in this manner, relative to balancer viability. Replicate crosses were performed and analyzed. Between 457 and 776 F1 animals (median = 647) were counted in the replicate *ago1* crosses. Between 234 and 380 F1 animals (median = 285) were counted in the replicate *dcr-1* crosses.

### Eye Mispatterning

Genetic mosaic animals bearing *mir-7^Δ1^* homozygous mutant eyes were generated using the FLP-FRT system. The animals’ genotype was: *w ey-FLP; FRT42D mir-7^Δ1^ / FRT42D GMR-Hid cl*. Matching wildtype control animals’ genotype was: *w ey-FLP; FRT42D P[w^+^] / FRT42D GMR-Hid cl*. Individuals also contained either *ILP2-GAL4* alone (control) or *ILP2-GAL4 UAS-Rpr* (IPC ablated) transgenes. All individuals were raised at 29°C. Eye roughening was scored as previously described (Li et al., 2009). For *RpS3* interactions with *mir-7*, trans-heterozygous *mir-7* mutants and matched wildtype controls (*Df(2R)exu1/+*) were raised at 29°C to adulthood. The *RpS3^2^* allele was combined with *mir-7* alleles. Eye roughening was scored as previously described (Li et al., 2009). Genetic mosaic animals bearing *ago1^W894^* homozygous mutant eyes were generated using the FLP-FRT system. The animals’ genotype was: *w ey-FLP; FRT42D ago1^W894^ / FRT42D GMR-Hid cl*. Matching wildtype control animals’ genotype was: *w ey-FLP; FRT42D P[w^+^] / FRT42D GMR-Hid cl*. Individuals also contained either *ILP2-GAL4* alone (control) or *ILP2-GAL4 UAS-Rpr* (IPC ablated) transgenes. For experiments with Yan transgenics, animals bearing one copy of either the *Yan^ACT^* or *Yan^WT^* (Rebay and Rubin, 1995) transgene also contained either *ILP2-GAL4* alone (control) or *ILP2-GAL4 UAS-Rpr* (IPC ablated) transgenes.

### Bristle Scoring

Animals of the correct genotype were allowed to age for 3 days after eclosion. The number of scutellar bristles was counted for each individual. Since these large bristles are positioned with high regularity and number on the scutellum, there was no ambiguity in counting the scutellar bristle number. For *wg* experiments, the number of sternopleural bristles was counted for each individual. Again, the position and number of these bristles is highly regular.

### Quantification of sfGFP-Sens Expression in the Wing Disc

Wing discs from white-prepupal females were dissected out in ice-cold Phosphate Buffered Saline (PBS). Discs were fixed in 4% paraformaldehyde in PBS for 20 minutes at 25°C and washed with PBS containing 0.3% Tween-20. Then they were stained with 0.5 μg/ml 4′,6-diamidino-2-phenylindole (DAPI) and mounted in Vectashield. Discs were mounted apical side up and imaged with identical settings using a Leica TCS SP5 confocal microscope. All images were acquired at 100x magnification at 2048 x 2048 resolution with a 75 nm x-y pixel size and 0.42 μm z separation. Scans were collected bidirectionally at 400 MHz and 6x line averaged. Wing discs of different genotypes were mounted on the same microscope slide and imaged in the same session for consistency in data quality.

For each wing disc, five optical slices containing Sens-positive cells along the anterior wing margin were chosen for imaging and analysis. A previously documented custom MATLAB script was used to segment nuclei in each slice of the DAPI channel (Pelaez et al., 2015). High intensity nucleolar spots were smoothed out to merge with the nuclear area to prevent spurious segmentation. Next, cell nuclei were identified by thresholding based on DAPI channel intensity. Segmentation parameters were optimized to obtain nuclei with at least 100 pixels and no more than 4000 pixels.

The majority of cells imaged did not reside within the proneural region and therefore displayed background levels of fluorescence scattered around some mean level. We calculated the “mean background” in the green channel of each disc individually. We did this by fitting a Gaussian distribution to the population and finding the mean of that fit. In order to separate sfGFP-Sens-positive cells, we chose a cut-off percentile based on the normal distribution, below which cells were deemed sfGFP-Sens-negative. We set this cut-off at the 84th percentile for all analysis since empirically it provided the most accurate identification of proneural cells (Giri et al., 2019). To normalize measurements across tissues and experiments, this value was subtracted from the total measured fluorescence for all cells in that disc. Only cells with values above the threshold for sfGFP fluorescence were assumed Sens positive (usually 30% of total cells) and carried forward for further analysis.

We analyzed >10 replicate wing discs for each treatment. In total, we measured wildtype *sfGFP-Sens* expression in 50,788 cells from wildtype *RpS13* discs and 18,945 cells from discs heterozygous mutant for *RpS13^1^*. We measured mutant *sfGFP-Sens^m1m2^* expression in 79,835 cells from wildtype *RpS13* discs and 12,954 cells from discs heterozygous mutant for *RpS13^1^*. The probability density plots were bootstrapped 1000 times by sampling with replacement, and the 95% CI was calculated by computing the 2.5/97.5th percentile y-value of probability density for each x-value.

### Quantification of Yan-YFP Expression Dynamics in the Eye

White-prepupal eye discs were dissected, fixed, and imaged by confocal microscopy for YFP and RFP fluorescence, as previously described (Pelaez et al., 2015). Briefly, samples fixed in 4% paraformaldehyde were kept in the dark at −20°C and imaged no later than 18-24 hr after fixation. In all cases, 1024 x 1024 16-bit images were captured using a Leica SP5 confocal microscope equipped with 40X oil objective. During imaging, discs were oriented with the equator parallel to the x-axis of the image. Optical slices were set at 0.8µm slices (45-60 optical slices) with an additional digital zoom of 1.2-1.4 to completely image eye discs from basal to apical surfaces. Images recorded a region of at least 6 rows of ommatidia on each side or the dorsal-ventral eye disc equator. All discs for a given condition were fixed, mounted, and imaged in parallel to reduce measurement error. Sample preparation, imaging, and analysis were not performed under blind conditions. Image data was processed for automatic segmentation and quantitation of RFP and YFP nuclear fluorescence as described (Pelaez et al., 2015). Briefly, cell segmentation was performed using a H2Av-mRFP marker as a reference channel for identification of cell nuclei boundaries. Each layer of the reference channel was segmented independently. A single contour containing each unique cell was manually selected and assigned a cell type using a custom graphic user interface. For each annotated cell contour, expression measurements were obtained by normalizing the mean pixel fluorescence of the YFP channel by the mean fluorescence of the His-RFP channel. This normalization serves to mitigate variability due to potentially uneven sample illumination, segment area, and differences in protein expression capacity between cells. We assigned cell-type identities to segmented nuclei by using nuclear position and morphology, two key features that enable one to unambiguously identify eye cell types without the need for cell-specific markers (Wolff and Ready, 1993). This task was accomplished using *FlyEye Silhouette*; an open-source package for macOS that integrates our image segmentation algorithm with a GUI for cell type annotation. Subsequent analysis and visualization procedures were implemented in Python.

Cell positions along the anterior-posterior axis were mapped to developmental time as described previously (Pelaez et al., 2015). This depends on two assumptions that have been extensively validated in the literature. One, the furrow proceeds at a constant velocity of one column of R8 neurons per two hours, and two, minimal cell migration occurs. For each disc, Delaunay triangulations were used to estimate the median distance between adjacent columns of R8 neurons. Dividing the furrow velocity by the median distance yields a single conversion factor from position along the anterior-posterior axis to developmental time. This factor was applied to all cell measurements within the corresponding disc. This method does not measure single cell dynamics, but rather aggregate dynamics across the developmental time course of cells in the eye.

Moving averages were computed by evaluating the median value among a collection of point estimates for the mean generated within a sliding time window. Each point estimate was generated via a hierarchical bootstrapping technique in which we resampled the set of eye discs, then resampled the aggregate pool of cell measurements between them. This novel method enhances our existing approach (Pelaez et al., 2015) by capturing variation due to the discretized nature of eye disc sample collection. Using the existing method, the error bars are considerably narrower (not shown). A window size of 500 sequential progenitor cells was used in all cases, but our conclusions are not sensitive to our choice of window size. Yan level measurements were pooled across multiple replicate eye discs. An automated approach was used to align these replicate samples in time. First, a disc was randomly chosen to serve as the reference population for the alignment of all subsequent replicates. Cells from each replicate disc were then aligned with the reference population by shifting them in time. The magnitude of this shift was determined by maximizing the cross-correlation of moving-average Yan-YFP expression *Y(t)* with the corresponding reference time series *X(t)*. Rather than raw measurements, moving averages within a window of ten cells were used to improve robustness against noise. This operation amounts to:

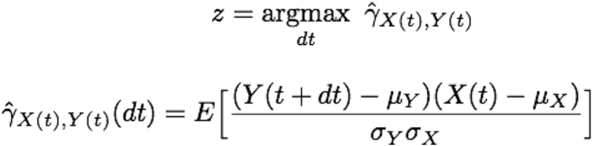

where, *μ* and *σ* are the mean and standard deviation of each time series, and *dt* is the magnitude of the time shift.

Different experimental treatments (e.g. wildtype and miR-7 null) were aligned by first aligning the discs within each treatment, then aggregating all cells within each treatment and repeating the procedure with the first treatment serving as the reference. We analyzed four to seven replicate eye discs for each treatment in two separate experiments. In total, we measured wildtype *Yan-YFP* levels in 4,518 cells in normally metabolizing samples and 4,379 cells in slowly metabolizing samples. We measured mutant *Yan^ΔmiR-7^-YFP* levels in 5,382 cells in normally metabolizing samples and 6,716 cells in slowly metabolizing samples.

### Mathematical Modeling Framework

Our model serves to demonstrate a general feature of dynamic systems subject to multiple auxiliary modes of feedback rather than reproducing specific regulatory mechanisms behind each experimental observation. We employ a control theoretic description of the transiently-induced expression and regulation of a single gene within an intracellular cascade of developmental gene expression. In control terminology, protein level is maintained at a basal steady state by proportional feedback in response to a disturbance. The disturbance induces activation of a gene, which induces transcription of mRNAs, which proceed to induce translation of protein. This corresponds to three sequential first-order transfer functions with interspersed feedback relating deviations in the input to deviations in output protein level (Fig. S1A).

A linear time invariant system describes the time evolution of activated DNA (*D*), mRNA (*R*), and protein (*P*) state variables in response to a disturbance (*I*) inducing gene activation. These discrete state variables describe the extent of gene expression at any point in time. Transitions between each of the variables’ states are governed by the set of linear reaction propensities listed below.

| Reaction | State Transition | Propensity | Parameter Value [$\text{min}^{-1}$] |
| --- | --- | --- | --- |
| gene activation | $\Delta D \rightarrow \Delta D + 1$ | $k_1 \Delta I$ | 1 |
| transcription | $\Delta R \rightarrow \Delta R + 1$ | $k_2 \Delta D$ | 1 |
| translation | $\Delta P \rightarrow \Delta P + 1$ | $k_3 \Delta R$ | 1 |
| gene deactivation | $\Delta D \rightarrow \Delta D - 1$ | $\gamma_1 \Delta D$ | 1 |
| transcript decay | $\Delta R \rightarrow \Delta R - 1$ | $\gamma_2 \Delta R$ | $1 \times 10^{-2}$ |
| protein decay | $\Delta P \rightarrow \Delta P - 1$ | $\gamma_3 \Delta P$ | $1 \times 10^{-3}$ |
| transcriptional feedback | $\Delta D \rightarrow \Delta D - 1$ | $\eta_1 \Delta P$ | $5.0 \times 10^{-4}$ |
| post-transcriptional feedback | $\Delta R \rightarrow \Delta R - 1$ | $\eta_2 \Delta P$ | $1.0 \times 10^{-4}$ |
| post-translational feedback | $\Delta P \rightarrow \Delta P - 1$ | $\eta_3 \Delta P$ | $5.0 \times 10^{-4}$ |

Rather than explicitly defined regulatory mechanisms, we abstract all modes of regulation as independent linear feedback terms:

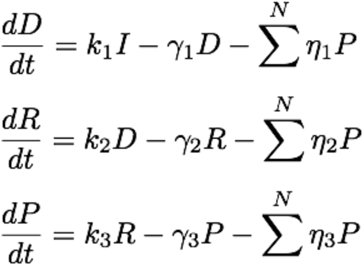

where *k_i_* are activation, transcription, or translation rate constants, *γ_i_* are degradation constants, *η_i_* are feedback strengths, and each species may be subject to *N* independent repressors. State variables are linearized about a steady state of zero stimulus:

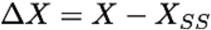

When expressed in the Laplace frequency domain this system consists of three algebraic equations relating deviations in input level to deviations in protein level:

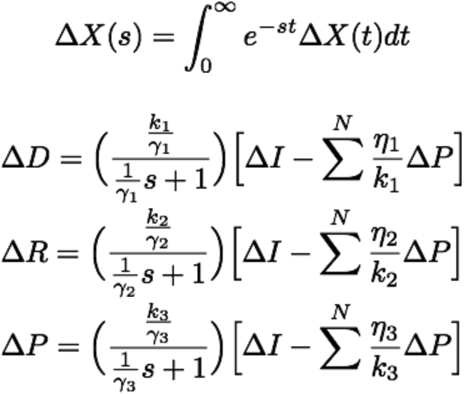

We represent gene activation, deactivation, transcription, transcript decay, translation, protein decay, and all modes of regulation using linear rate laws. Linear kinetics enables characterization of the transient-pulse system using a small number of parameters. Although protein synthesis and gene-product decay are typically modeled as linear processes, transcriptional and regulatory kinetics are frequently described by nonlinear propensities. Therefore, we also considered two nonlinear modeling frameworks, both of which recapitulated the results using linear kinetics (Figs. S2F,G and S4F,G).

### Dependence of Model Parameters on Metabolic Conditions

IPC ablation reduces cellular glucose consumption. Presumably this would affect either the production and consumption of ATP or the production and consumption of substrates for RNA and protein synthesis (or both). The precise effects are unknown, so we independently modeled each scenario. Since ATP concentration remains fairly constant when respiration is limited (Brown, 1992), ATP flux (and ATP synthesis) is assumed to decrease. Because transcription, translation, and protein degradation all require ATP turnover, we halved their rate parameters under conditions of reduced glucose consumption. Under conditions of reduced substrate availability for RNA/protein synthesis, we assumed that only transcription and translation rates are affected by limiting fluxes of nucleotides and amino acids. We assumed only the translation rate is affected under conditions of reduced ribosome number. These assumptions are incorporated as changes to the model’s rate parameters as listed below.

| Parameter | Normal Metabolism | Reduced ATP Consumption | Reduced RNA/ Protein Substrates | Reduced Ribosome Number |
| --- | --- | --- | --- | --- |
| transcription rate constant | $k_2$ | $\frac{1}{2} k_2$ | $\frac{1}{2} k_2$ | $k_2$ |
| translation rate constant | $k_3$ | $\frac{1}{2} k_3$ | $\frac{1}{2} k_3$ | $\frac{1}{2} k_3$ |
| protein decay rate constant | $\gamma_3$ | $\frac{1}{2} \gamma_3$ | $\gamma_3$ | $\gamma_3$ |

In all cases, feedback strengths were reduced in order to account for the intermediate processes abstracted by each feedback element. Feedback strength parameters (*η_i_*) were reduced four-fold under conditions of reduced energy metabolism and reduced RNA/protein substrate availability. This scaling assumes that both transcription and translation occur within the arbitrarily complex regulatory motifs represented by each repressor. This is a reasonable assumption for repressor proteins such as transcription factors and kinases. For RNA repressors such as microRNAs, feedback strength parameters could instead be reduced only two-fold to account for their reduced transcription rates. However, microRNAs must be transcribed, processed, and act with effector proteins in order to repress their targets. These fourfold reductions in feedback strength correspond to fourfold reduction of the transcriptional feedback gain (*K_C1_*) and twofold reduction in the post-transcriptional and post-translational feedback gains (*K_C2_* and *K_C3_*). Feedback strength parameters (*η_i_*) were only reduced two-fold under reduced protein synthesis conditions. This implies that the transcriptional and post-transcriptional feedback gains (*K_C1_* and *K_C2_*) decrease twofold while the post-translational feedback gain (*K_C3_*) remains constant. Each of these dependencies are summarized below.

| Parameter | Normal Metabolism | Reduced ATP Consumption | Reduced RNA/ Protein Substrates | Reduced Ribosome Number |
| --- | --- | --- | --- | --- |
| transcriptional feedback strength | $\eta_1$ | $\frac{1}{4}\eta_1$ | $\frac{1}{4}\eta_1$ | $\frac{1}{2}\eta_1$ |
| post-transcriptional feedback strength | $\eta_2$ | $\frac{1}{4}\eta_2$ | $\frac{1}{4}\eta_2$ | $\frac{1}{2}\eta_2$ |
| post-translational feedback strength | $\eta_3$ | $\frac{1}{4}\eta_3$ | $\frac{1}{4}\eta_3$ | $\frac{1}{2}\eta_3$ |

### Model Simulations

Default parameter values were based on approximate transcript and protein synthesis and turnover rates for animal cells reported in the literature (Milo and Phillips, 2016), while gene activation and decay rates were arbitrarily set to a significantly faster timescale. Default feedback strengths for repressors acting at the gene, transcript, or protein levels were chosen such that ~25-50% of simulations failed to reach the threshold under normal conditions when one of two identical repressors was lost. Population-wide expression dynamics were estimated by simulating 5000 output trajectories in response to a three-hour transient step input to the gene activation rate. Simulations were performed using a custom implementation of the stochastic simulation algorithm (Gillespie, 1977). The algorithm constrains solutions to the set of discrete positive values, consistent with linearization about a basal level of zero gene activity. This simplifying assumption is based on the near-zero basal activities expected in the experimental systems, but is not required to support the conclusions of the model (Figs. S2D and S4D).

### Evaluation of Error Frequencies and Changes in Expression Dynamics

Gene expression trajectories were simulated both with (full repression) and without (partial repression) a second repressor. The time point at which the full-repression simulations mean level reached 30% of its maximum value was taken to be the commitment time. At this time, a threshold for developmental success was set at the 99^th^ percentile of protein levels subject to full-repression. Error frequencies were obtained by evaluating the fraction of simulated protein levels that exceeded this threshold. Per this definition, the minimum possible error frequency is one percent. For simplicity we subtracted this percentage point from all reported error frequencies.

Protein expression dynamics were compared by evaluating the fraction of partially-repressed simulation trajectories in excess of the 99^th^ percentile of fully-repressed trajectories at each point in time. These fractions were then averaged across the time course, beginning with the reception of the input signal and ending at the previously defined commitment time. Each fraction may be thought of as the instantaneous error frequency, and their average reflects the extent to which the expression dynamics differ between the two sets of simulated trajectories (Figs. S3A,B).

### Parameter Variation and Sensitivity to Model Assumptions

We conducted a systematic parameter sweep in which all parameters were varied across a ten-fold range (± ~three-fold). For each parameter set we ran six sets of five thousand simulations: 1) full feedback with normal metabolism and translation, 2) partial feedback with normal metabolism and translation, 3) full feedback with reduced energy metabolism, 4) partial feedback with reduced energy metabolism, 5) full feedback with reduced protein synthesis, 6) partial feedback with reduced protein synthesis. Full-repression systems were assigned two copies of each feedback element present in the corresponding partial-repression system. Error frequencies were evaluated as described above. This procedure constitutes one parameter sweep.

Error frequency is greater than 1% for almost all combinations of parameter values (Figs. S1B,C), indicating that partial loss of repression induces an increase in error frequency across a broad parameter range. We also varied the level of the success threshold, and recalculated all error frequencies accordingly. Error frequency is greater than 1% for almost all definitions of the success threshold, indicating that loss of a repressor increases developmental error irrespective of where the success threshold is set (Fig. S1D).

The differences in error frequency between simulations with normal metabolism and reduced metabolism are shown in Fig. S2B for all parameter sets, while the corresponding difference between simulations with normal protein synthesis and reduced protein synthesis are shown in Fig. S4A. There is a general trend of decreased error frequency with partial feedback under both reduced energy metabolism and reduced protein synthesis conditions, irrespective of where the success threshold is set (Figs. S2C and S4C).

Our conclusion also persists when a nonzero basal stimulus is introduced. We conducted an additional parameter sweep in which the stimulus consists of a transient step change between input values of Δ*I*=0.1 and Δ*I* =1.0. Simulations were carried out on an absolute basis, and were allowed sufficient time to reach a non-zero steady state before and after the stimulus was applied. The resultant protein level trajectories for each of the six sets of simulations were converted to deviation form by subtracting the respective population-wide mean final value. Error frequencies were then evaluated as previously described. Despite the inclusion of a nonzero basal stimulus, error frequencies remained broadly suppressed under conditions of both reduced energy metabolism and reduced protein synthesis (Figs. S2D and S4D).

The preceding simulations assume the stimulus (input) is a unit step that persists for three hours regardless of metabolic conditions (Figs. S2B and S4A). Alternatively, metabolic conditions might affect stimulus (input) duration, particularly if the upstream processes responsible for the input are also governed by metabolically delayed processes. We find that the general prediction made by our model – that reduced energy metabolism and reduced protein synthesis limit sensitivity to loss of regulation – persists in roughly half of cases if we apply four-fold and two-fold extensions of input duration under reduced energy metabolism and reduced protein synthesis conditions, respectively (Figs. S2E and S4E). Notably, in many cases scaling the input duration with metabolic condition yields the opposite effect. However, these instances correspond to simulations in which the extended stimulus yields output protein levels greater than those observed under normal metabolic conditions, suggesting that a four-fold increase in stimulus duration may be excessive.

### Alternate Models

The number of active sites firing transcription within a cell is limited by gene copy number, but the activated-DNA state in our simple linear model is unbounded. To test whether error frequency suppression persists when an upper bound on gene activity is introduced, we considered a simple two-state transcription model

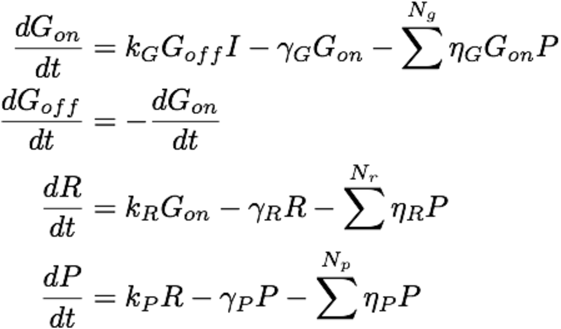

where *G_on_* and *G_off_* are the on- and off-states of a gene; *I*, *R* and *P* are the input, transcript, and protein levels; *k_i_*, *γ*_*i*_, and *η*_*i*_ are the synthesis, decay, and feedback rate constants for species *i*; and *N_g_*, *N_r_*, and *N_p_* are the number of transcriptional, post-transcriptional, and post-translational repressors, respectively. Rate parameter dependencies upon metabolic and protein synthesis conditions were analogous to those used in the linear model, and are tabulated below.

| Parameter | Normal Metabolism | Reduced Metabolism | Reduced Protein Synthesis |
| --- | --- | --- | --- |
| transcription rate constant | $k_R$ | $\frac{1}{2} k_R$ | $k_R$ |
| translation rate constant | $k_P$ | $\frac{1}{2} k_P$ | $\frac{1}{2} k_P$ |
| protein decay rate constant | $\gamma_P$ | $\frac{1}{2} \gamma_P$ | $\gamma_P$ |
| transcriptional feedback strength | $\eta_G$ | $\frac{1}{4} \eta_G$ | $\frac{1}{2} \eta_G$ |
| post-transcriptional feedback strength | $\eta_R$ | $\frac{1}{4} \eta_R$ | $\frac{1}{2} \eta_R$ |
| post-translational feedback strength | $\eta_P$ | $\frac{1}{4} \eta_P$ | $\frac{1}{2} \eta_P$ |

We performed another parameter sweep varying each of the model’s nine parameters across one order of magnitude. All simulations were initialized as diploid (*G_off_* = 2) then subject to a constant 3-hour stimulus before reverting to a basal level of zero gene expression. Despite the limitation placed on gene activity, error frequency remains elevated under normal growth conditions and broadly suppressed when metabolism or protein synthesis are reduced (Figs. S2F and S4F).

Gene expression models also frequently utilize cooperative kinetics in order to capture the nonlinearities and thresholds encountered in transcriptional regulation. We reformulated our gene expression model in terms of Hill kinetics:

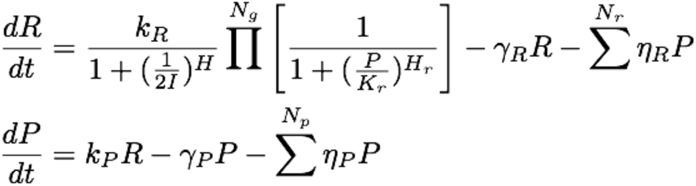

where *I*, *R*, and *P* are the input, transcript, and protein levels; *k_i_*, γ*i*, and η*i* are the synthesis, decay, and linear feedback rate constants for species *i*; *N_r_*, and *N_p_* are the number of post-transcriptional, and post-translational linear repressors; *H* is a transcriptional Hill coefficient; and *K_r_* and *H_r_* are the half-maximal occupancy level and Hill coefficient of each of the *N_g_* transcriptional repressors. The stimulus level corresponding to half-maximal transcription rate was fixed at 0.5 because we only consider a binary input signal. Rate parameters were again scaled with metabolic and protein synthesis conditions in a manner analogous to the linear model.

| Parameter | Normal Metabolism | Reduced Metabolism | Reduced Protein Synthesis |
| --- | --- | --- | --- |
| transcription rate constant | $k_R$ | $\frac{1}{2} k_R$ | $k_R$ |
| translation rate constant | $k_P$ | $\frac{1}{2} k_P$ | $\frac{1}{2} k_P$ |
| protein decay rate constant | $\gamma_P$ | $\frac{1}{2} \gamma_P$ | $\gamma_P$ |
| post-transcriptional feedback strength | $\eta_R$ | $\frac{1}{4} \eta_R$ | $\frac{1}{2} \eta_R$ |
| post-translational feedback strength | $\eta_P$ | $\frac{1}{4} \eta_P$ | $\frac{1}{2} \eta_P$ |

The half-maximal occupancy level and Hill coefficients of transcriptional repressors were assumed to be independent of growth rate. Another parameter sweep revealed that despite the incorporation of cooperative binding kinetics, error frequency remains elevated under normal metabolic conditions and is broadly suppressed when metabolism or protein synthesis are reduced (Figs. S2G and S4G).

### QUANTIFICATION AND STATISTICAL ANALYSIS

Population proportions were compared using a Chi-square test with Yates’ correction and a Fisher’s exact test. Both tests gave similar results. All tests involving multiple experimental groups were Bonferroni corrected. Analysis of *sev* experiments scoring R7 cells applied a one-way ANOVA with Bonferroni correction. Relative viabilities were compared using a Mann-Whitney-Wilcoxon test with Bonferroni correction. These tests were performed using Prism 7 (GraphPad) software. P-values shown in figures are presented from tests with the most conservative value shown if more than one test was performed on data. * *p* < 0.05; ** *p* < 0.01; *** *p* < 0.001; **** *p* < 0.0001 Confidence intervals for the moving average of Yan-YFP expression were inferred from the 2.5th and 97.5th percentile of 1000 point estimates of the mean within each moving-average window. Point estimates were generated by bootstrap resampling with replacement of the expression levels within each window.

Analysis of sfGFP-Sens fluorescence was performed using two independent approaches. 1) For each genotype, 1000 point-estimates were made of the median fluorescence level in cells. Point estimates were generated by bootstrap resampling with replacement of the cell samples within each genotype. Point estimates from wildtype sfGFP-Sens were then randomly paired with point estimates from miR-9a-resistant sfGFP-Sens to derive a set of 1000 point-estimates of the fold-change in median sfGFP-Sens expression. Confidence intervals for the average fold-change in sfGFP-Sens expression were inferred from the 0.5th and 99.5th percentile of these point estimates. 2) The distributions of fluorescence from wildtype *sfGFP-sens* and mutant *sfGFP-sens^m1m2^* cell populations were compared using a Mann-Whitney-Wilcoxon test implemented in R. By calculating the difference between all randomly paired cell samples from wildtype versus mutant, the location shift μ is estimated as the median of the difference between a sample from sfGFP-Sens and a sample from sfGFP-Sensm1m2. Confidence intervals for the shift were inferred from the 2.5th and 97.5th percentile of the set of differences.

There was no exclusion of any data or subjects.

---

**Figure S1.**
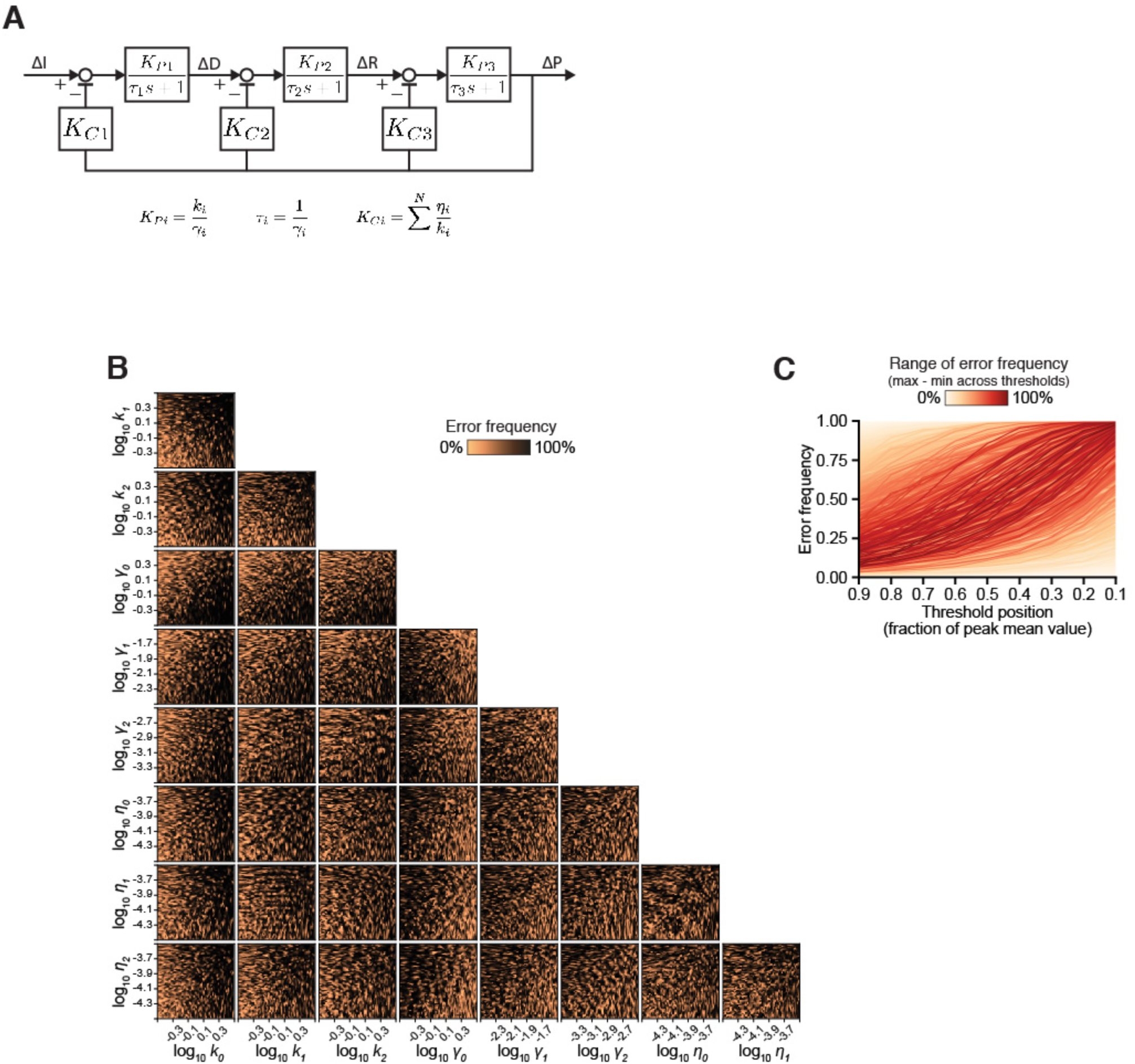
Related to Figure 4. Model demonstrates that error frequencies are broadly increased when an auxiliary repressor is lost. **(A)** Block diagram depiction of the mathematical model. Boxes contain transfer functions relating upstream and downstream variables, open circles indicate summation points. Transfer functions are expressed in the Laplace frequency domain. **(B)** Simulations were performed with 2500 independent parameter sets quasi-randomly sampled from the nine-dimensional hyperspace defined by one order of magnitude variation in each of the model parameters. Error frequencies are projected onto two dimensional planes and then linearly interpolated onto (100 px)^2^ grids. **(C)** Error frequencies for parameter sets from (B) were re-calculated across a range of different success thresholds. Shown are error frequencies incurred by the loss of one repressor. Thresholds are defined by the 99^th^ percentile of protein levels simulated with all repressors, and are evaluated at the time when the mean protein level with all repressors reaches the indicated fraction of its maximum value. Each line represents one of the parameter sets from (B). Color scale reflects maximum difference in error frequency across the range of thresholds tested.

**Figure S2.**
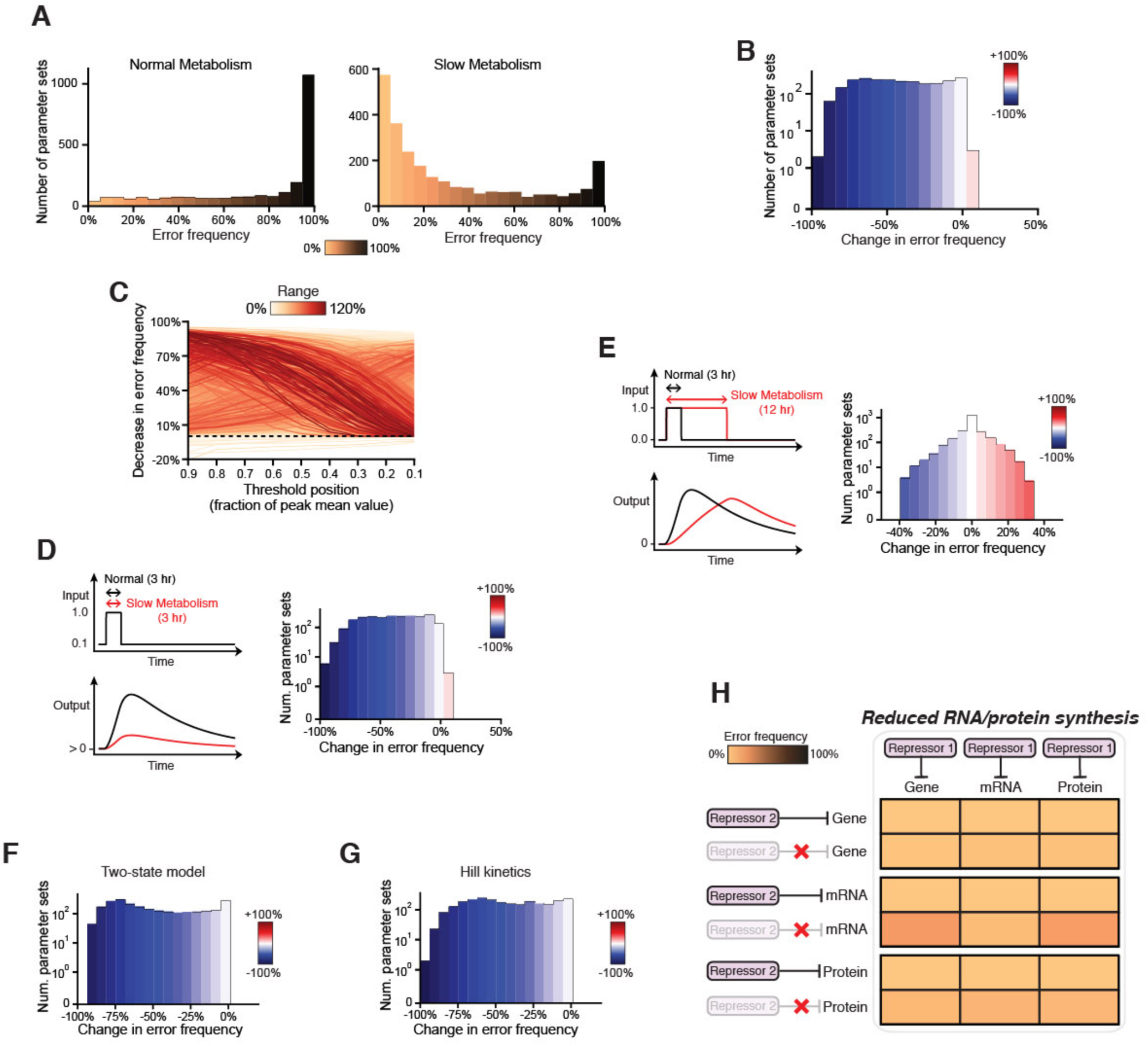
Related to Figure 4. Reduced energy metabolism diminishes the importance of auxiliary repressors over a wide range of model conditions. **(A-G)** For each panel, 2500 simulations were performed with parameter sets quasi-randomly sampled from the nine-dimensional hyperspace defined by one order of magnitude variation in each of the respective model parameters, as done for Fig. S1B. For each parameter set, error frequencies pertain to 50% loss of repression mimicking partial repressor loss. **(A)** One-dimensional representation of the error frequencies for all parameter sets under conditions of normal or diminished energy metabolism. **(B)** Change in error frequencies with diminished metabolism relative to normal metabolic conditions for all parameter sets. Blue-Red color scale corresponds to the difference in error frequency between low-metabolic and normal conditions, e.g. blue indicates error suppression by reduced energy metabolism. **(C)** Results from (B) re-calculated across a range of different success thresholds. Each line corresponds to a single parameter set. Color scale reflects maximum change in error frequency across the threshold range. Black dashed line corresponds to unchanged error frequency by reduced energy metabolism. The vast majority of simulations exhibit some reduction in error frequency across all thresholds. **(D-G)** Systematic modification of model conditions showing the change in error frequencies with diminished metabolism relative to normal metabolic conditions for all parameter sets. Blue-Red color scale corresponds to the difference in error frequency between low-metabolic and normal conditions, e.g. blue indicates error suppression by reduced energy metabolism. **(D)** Simulations where a nonzero basal stimulus is applied. **(E)** Simulations where input duration is increased four-fold by a reduction in energy metabolism. **(F)** Simulations when an upper bound is placed on the number of sites firing transcription. **(G)** Simulations when cooperative transcription kinetics are considered. **(H)** Partial repression imparts few developmental errors when RNA and protein synthesis rate parameter values are reduced by 50%. Construction is otherwise analogous to the right panel of Figure 4G.

**Figure S3.**
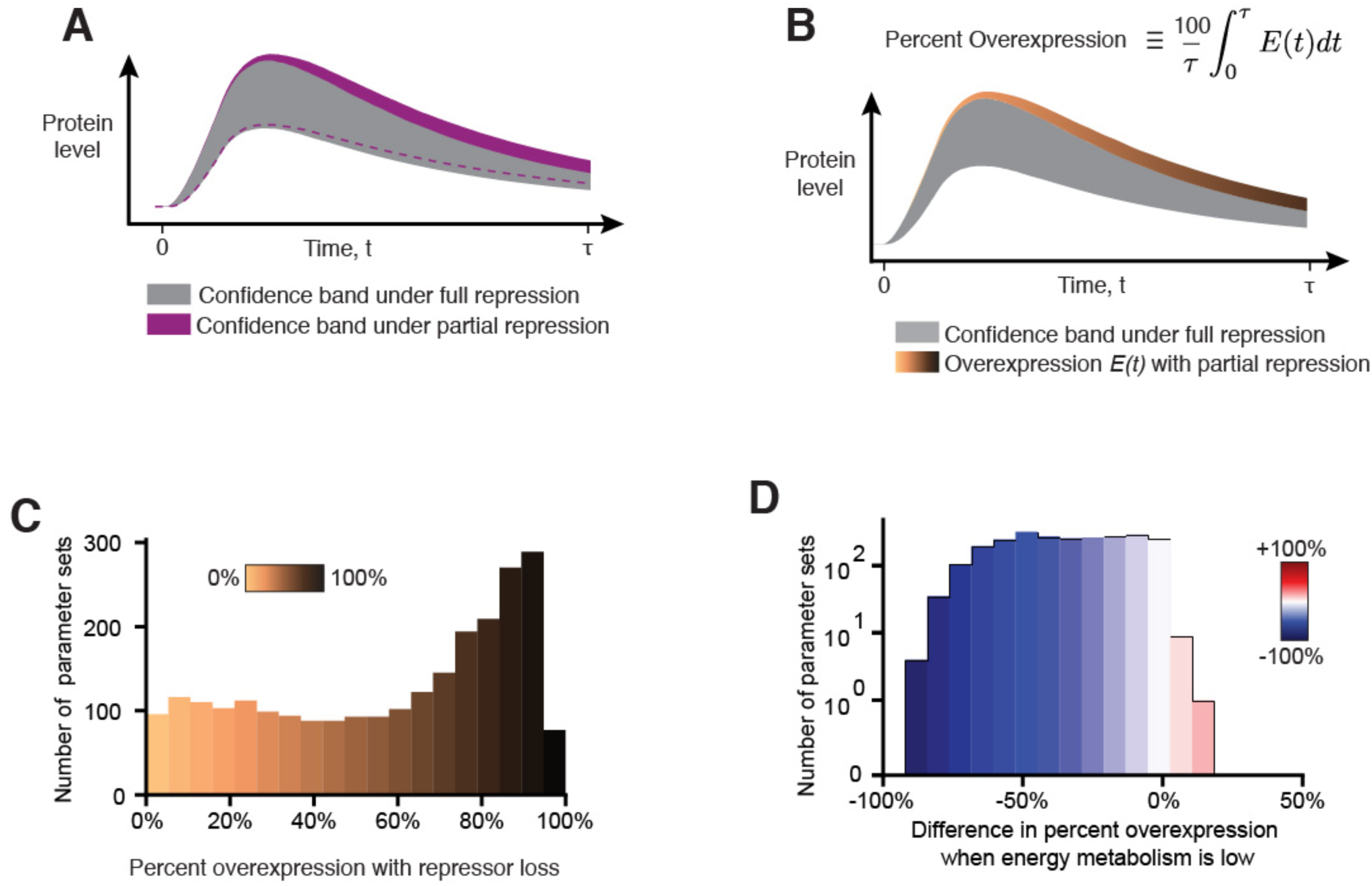
Related to Figure 5. Reductions in energy metabolism limit the extent to which protein expression dynamics are affected by loss of a repressor. **(A-B)** Graphical depiction of the method used to quantify the impact of repressor loss on protein expression dynamics. (A) Confidence bands span the 1st to 99^th^ quantiles of protein levels simulated under full repression (grey) and under partial repression (purple), where a repressor is lost. The dashed purple line denotes the lower bound of the purple confidence band. The symbol τ denotes the commitment time as defined previously. **(B)** Loss of a repressor causes protein overexpression E(t), which is calculated as the fraction of simulations that exceed the confidence band observed under full repression (grey) at a given time point. Orange-brown color scale reflects the value of E(t) for each time point in the time course. Percent overexpression is calculated as a percent of simulations that exceed the confidence band integrated over the entire time course. A maximum of 100% overexpression would occur when all simulations exceed the confidence band at all timepoints. **(C)** Percent overexpression caused by loss of a repressor for model simulations performed with 2500 independent parameter sets. Color scale reflects the strength of overexpression. Overexpression is large for most parameter sets. **(D)** Percent overexpression caused by loss of a repressor was calculated for simulations implementing normal energy metabolism and reduced energy metabolism. The difference in percent overexpression between the two metabolic conditions is shown for model simulations performed with 2500 independent parameter sets. Color scale reflects the difference. The majority of simulations are blue, indicating that expression dynamics are less affected by repressor loss when energy metabolism is low.

**Figure S4.**
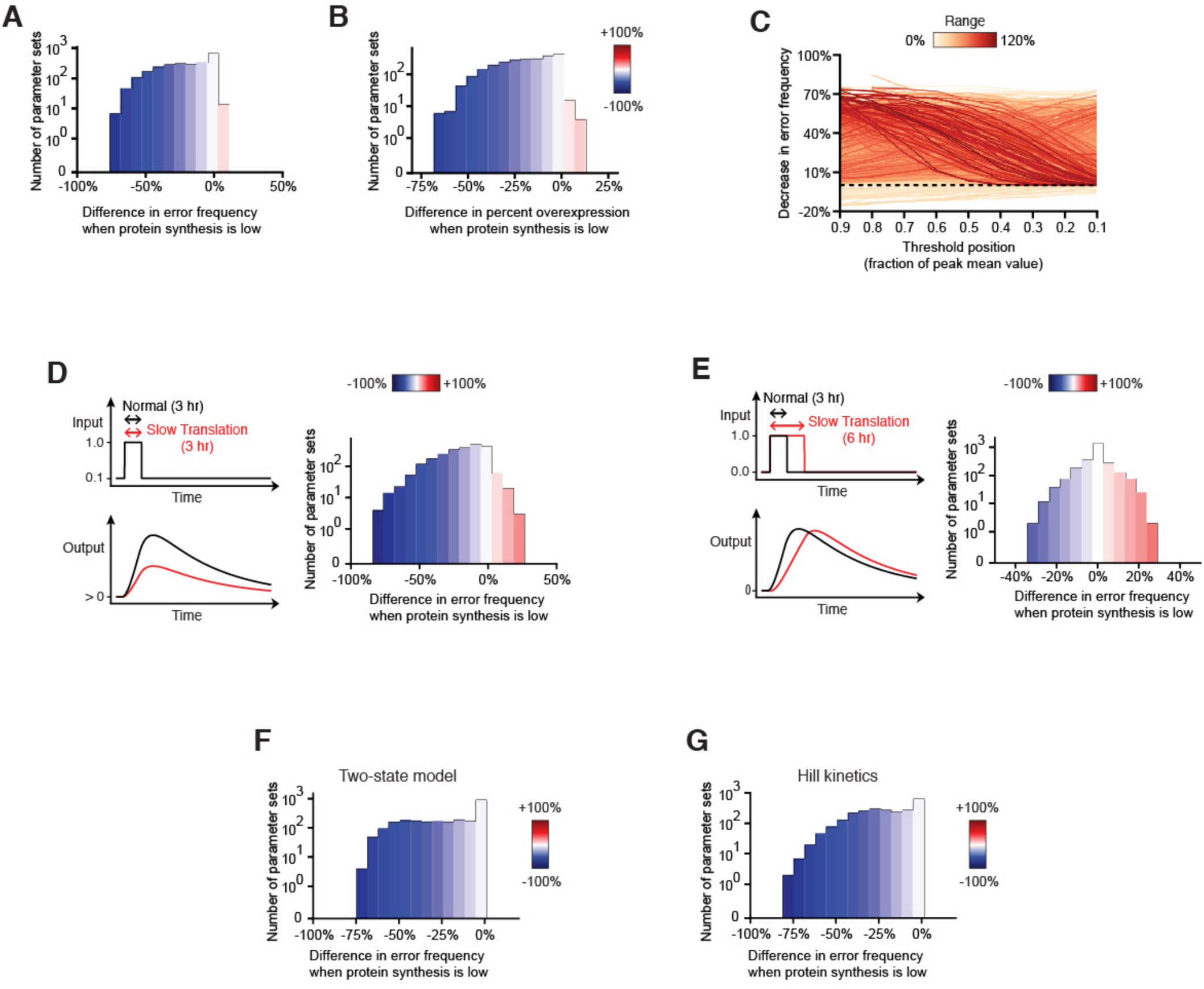
Related to Figure 7. Reduced protein synthesis capacity diminishes the importance of auxiliary repression over a wide range of model conditions. Each panel depicts a parameter sweep of the nine-dimensional hyperspace defined by one order of magnitude variation in each of the respective model parameters. For each parameter set, error frequency and percent overexpression were calculated as described earlier in Figs. S2 and S3. They pertain to 50% reduced repression mimicking auxiliary repressor loss. Error frequency and percent overexpression were calculated independently for conditions of normal and reduced protein synthesis. The difference in error frequency or overexpression between metabolic conditions are shown color-coded, e.g. blue indicates error suppression by reduced protein synthesis. **(A-C)** Simulations in which the duration of the stimulus input is constant. Shown are the **(A)** differential error frequencies and **(B)** differential changes in expression dynamics relative to normal protein synthesis conditions. **(C)** Differential error frequencies for varying definitions of the success threshold. Each line represents one parameter set, colored by the corresponding range of differential error frequencies. **(D)** Simulations where a nonzero basal stimulus is applied. **(E)** Simulations where input duration is increased two-fold by a reduction in protein synthesis capacity. **(F)** Simulations when an upper bound is placed on the number of sites firing transcription. **(G)** Simulations when cooperative transcription kinetics are considered.

